# The radiation of alopiine clausiliids in the Sicilian Channel (Central Mediterranean): phylogeny, patterns of morphological diversification and implications for taxonomy and conservation of *Muticaria* and *Lampedusa*

**DOI:** 10.1101/208348

**Authors:** V. Fiorentino, N. Salomone, P.J. Schembri, G. Manganelli, F. Giusti

**Affiliations:** University of Potsdam, Unit of Evolutionary Biology/Systematic Zoology Institute of Biochemistry and Biology Karl-Liebknecht-Str. 24-25, Potsdam, Germany; University of Siena, Department of Evolutive Biology, Via A. Moro, 2, 53100 Siena, Italy; University of Malta, Department of Biology, Msida MSD2080, Malta; University of Siena, Department of Environmental Sciences, Via P.A. Mattioli 4, 53100 Siena, Italy

## Abstract

The phylogeny, biogeography and taxonomy of the alopiine clausiliids of the Sicilian Channel, belonging to the genera *Lampedusa* and *Muticaria*, were investigated using morphological (shell characters and anatomy of the reproductive system) and genetic (sequencing of a fragment of the mitochondrial large ribosomal subunit 16S rRNA, and the nuclear internal transcriber spacer 1, ITS-1 rRNA) data. Classically, the genus *Lampedusa* includes three species: *L. imitatrix* and *L. melitensis* occurring in circumscribed localities in western Malta and on the islet of Filfla, and *L. lopadusae* on Lampedusa and Lampione. The genus *Muticaria* includes two species in southeastern Sicily (*M. siracusana* and *M. neuteboomi*) and one in the Maltese islands (*M. macrostoma*), which is usually subdivided into four entities based on shell characters (*macrostoma* on Gozo, Comino, Cominotto and central-eastern Malta; *mamotica* in southeastern Gozo; *oscitans* on Gozo and central-western Malta; *scalaris* in northwestern Malta). These have sometimes been considered as subspecies and sometimes as mere morphs.

The *Lampedusa* of Lampedusa and Lampione form a well distinct clade from those of the Maltese Islands. The population of Lampione islet is a genetically distinct geographic form that deserves formal taxonomic recognition (as *L. nodulosa* or *L. l. nodulosa*). The *Lampedusa* of Malta are morphologically distinct evolutionary lineages with high levels of genetic divergence and are confirmed as distinct species (*L. imitatrix* and *L. melitensis*).

The *Muticaria* constitute a clearly different monophyletic clade divided into three geographical lineages corresponding to the Sicilian, Maltese and Gozitan populations. The Sicilian *Muticaria* form two morphologically and genetically distinguishable subclades that may either be considered subspecies of a polytypic species, or two distinct species. The relationships of Maltese and Gozitan *Muticaria* are complex. Two of the three Maltese morphotypes resulted monophyletic (*oscitans* and *scalaris*) while the other was separated in two lineages (*macrostoma*); however this picture may be biased as only few samples of *macrostoma* were available to study. The Gozitan morphotypes (*macrostoma*, *mamotica* and *oscitans*) where resolved as polyphyletic but with clear molecular evidence of mixing in some cases, indicating possible relatively recent differentiation of the Gozitan *Muticaria* or repetitive secondary contacts between different morphotypes. Definitive taxonomic conclusions from these results are premature. Maltese *Muticaria* could be subdivided into three taxa according to morphological and molecular data (*M. macrostoma* or *M. m. macrostoma*, *M. oscitans* or *M. m. oscitans and M. scalaris* or *M. m scalaris*). Gozitan *Muticaria* could be considered a distinct polytypic species (for which the oldest available name is *Muticaria mamotica)* subdivided into subspecies showing a morphological range from *macrostoma*-like to *mamotica*-like and *oscitans* like.

Only the two Maltese species of *Lampedusa* are legally protected (by the European Union’s ‘Habitats Directive’ and Maltese national legislation). The present study has shown that the alopiine clausiliids of the Sicilian Channel constitute a number of genetically and/or morphologically distinct populations that represent important pools of genetic diversity, with, in some cases, a very circumscribed distribution. As such, these populations deserve legal protection and management. It is argued that without formal taxonomic designation, it would be difficult to extend international legal protection to some of the more threatened of these populations.

## Introduction

Archipelagos are ideal places for studying evolutionary biology; their limited geographical area makes them “paradigm systems” for understanding the origin of diversity (Mayr, 1963).

In the past century, archipelagos have received much attention from conservation biologists since most are inhabited by species-rich groups in fragile equilibria with their environment. Numerous studies have focussed on insular groups of snails, for example, *Partula* in the Indo-Pacific islands (Johnson, Murray & Clarke, 1993; Goodacre & Wade, 2001), *Polymita* in Cuba (Reyes-tur, Fernández & Suárez, 2001), *Napaeus* in the Canary archipelago (Alonso et al., 2006), and leptaxine hygromiids in the Azores islands (Van Riel et al., 2003; Jordaens et al., 2009).

Mediterranean archipelagos, such as the Pelagian and Maltese island groups, are also rich in endemic land snails, especially the alopiine clausiliids and the trochoideine hydgromiids (Giusti, Manganelli & Schembri, 1995). The former, a group of clausiliid snails mostly diversified in the Balkan peninsula and in Crete, occur in the Italian area with four genera: *Leucostigma* and *Medora* in the Apennines, and *Lampedusa* and *Muticaria* in south-eastern Sicily and the Maltese and Pelagian islands (Nordsieck, 2007) (Fig. 1).

**Figure 1.**
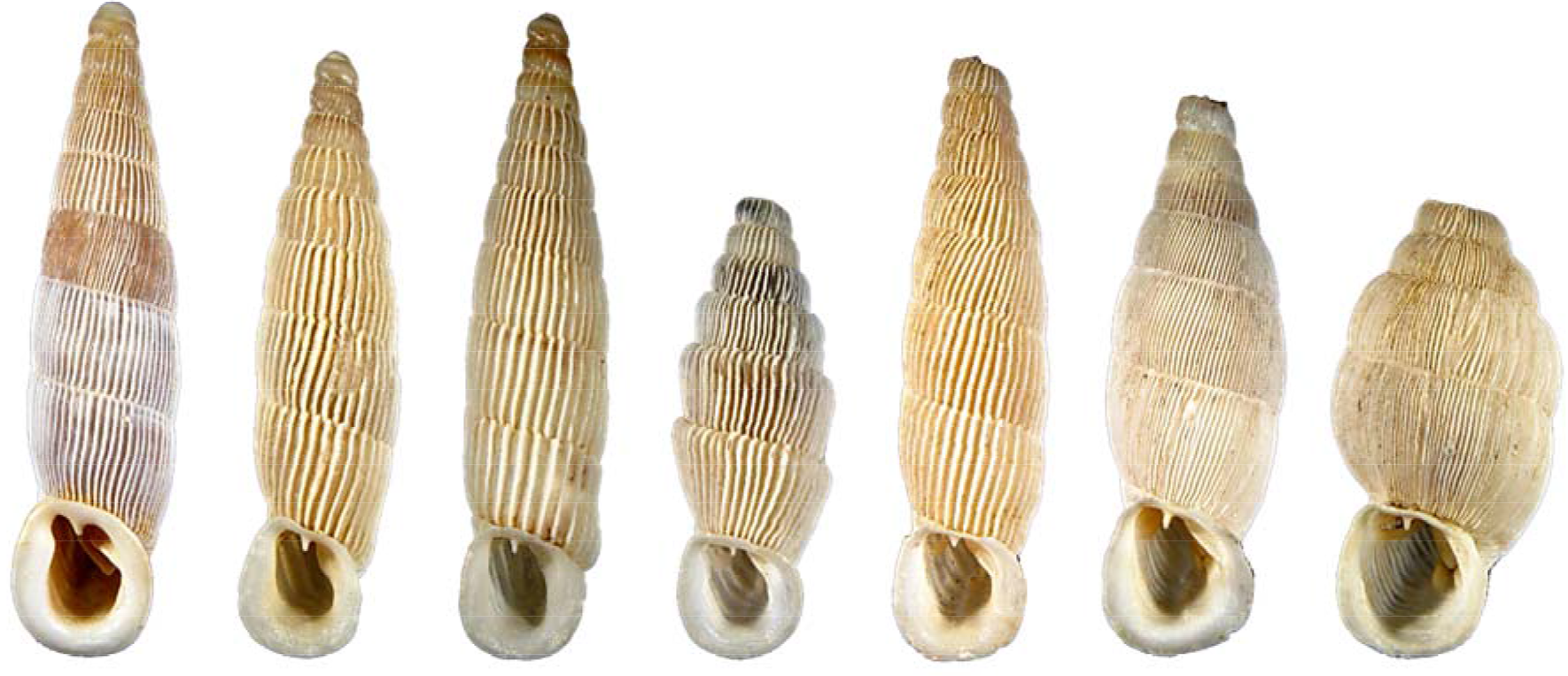
Representatives of the alopiine clausiliids of the Sicilian Channel: from left to right: *Lampedusa lopadusae*, *Lampedusa imitatrix*, *Lampedusa melitensis*, *Muticaria macrostoma* morph *scalaris, Muticaria macrostoma* morph *macrostoma, Muticaria macrostoma* morph *oscitans* and *Muticaria macrostoma* morph *mamotica*, according to current taxonomy (Giusti et al., 1995).

Most of the recognized taxa of *Lampedusa* and *Muticaria* exhibit allopatric or parapatric distribution patterns. The genus *Lampedusa* includes three species: *L. imitatrix* and *L. melitensis* occurring in two isolated areas in western Malta and on the islet of Filfla (5.2 km south of Malta), and *L. lopadusae* on Lampedusa and Lampione. Qualitative shell and anatomical characters distinguish *L. lopadusae* from *L. imitatrix* and *L. melitensis* (Giusti et al., 1995).

The genus *Muticaria* includes two species in south-eastern Sicily (*M siracusana* and *M. neuteboomi*) and one in the Maltese islands (*M. macrostoma*) (Fig. 2). Maltese *Muticaria* are usually subdivided into four entities on the base of shell shape and ribbing; these are sometimes considered as distinct species, sometimes as subspecies, and sometimes as morphs without taxonomic value (Table 1). The four are: *macrostoma* characterized by a conical fusiform shell with strong sparse ribs (Gozo, Comino, Cominotto and central-eastern Malta); *mamotica* characterized by a ventricose shell with slender minute close ribs (Xlendi Valley, near Munxar, southeastern Gozo); *oscitans* characterized by a conical fusiform shell with slender minute close ribs (Gozo and central-western Malta); and *scalaris* characterized by a conical scalariform shell with strong sparse ribs (Mistra Bay, northwestern Malta). Qualitative shell characters distinguish the Maltese *Muticaria* from the Sicilian species but no anatomical feature seems to distinguish the different Maltese forms (Giusti et al., 1995).

**Table 1.**
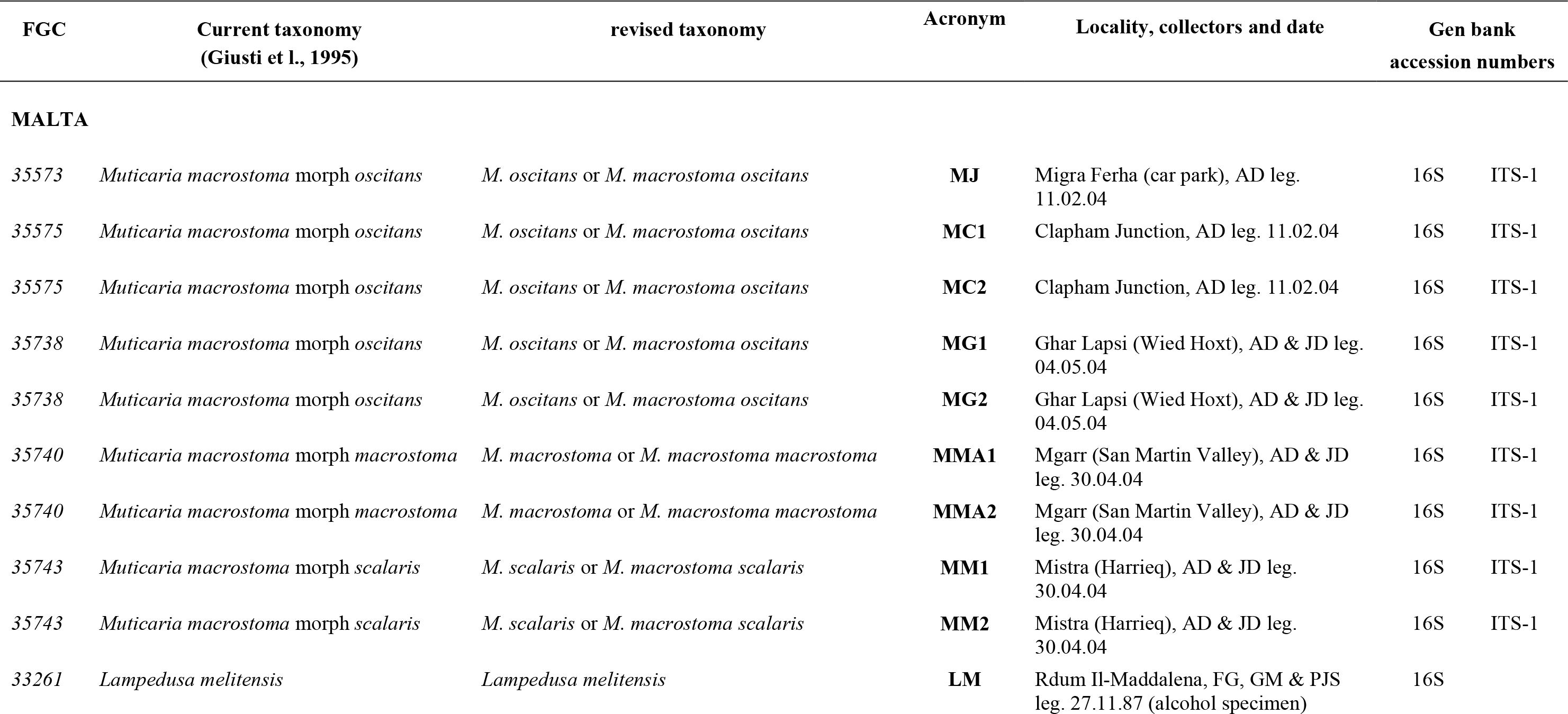
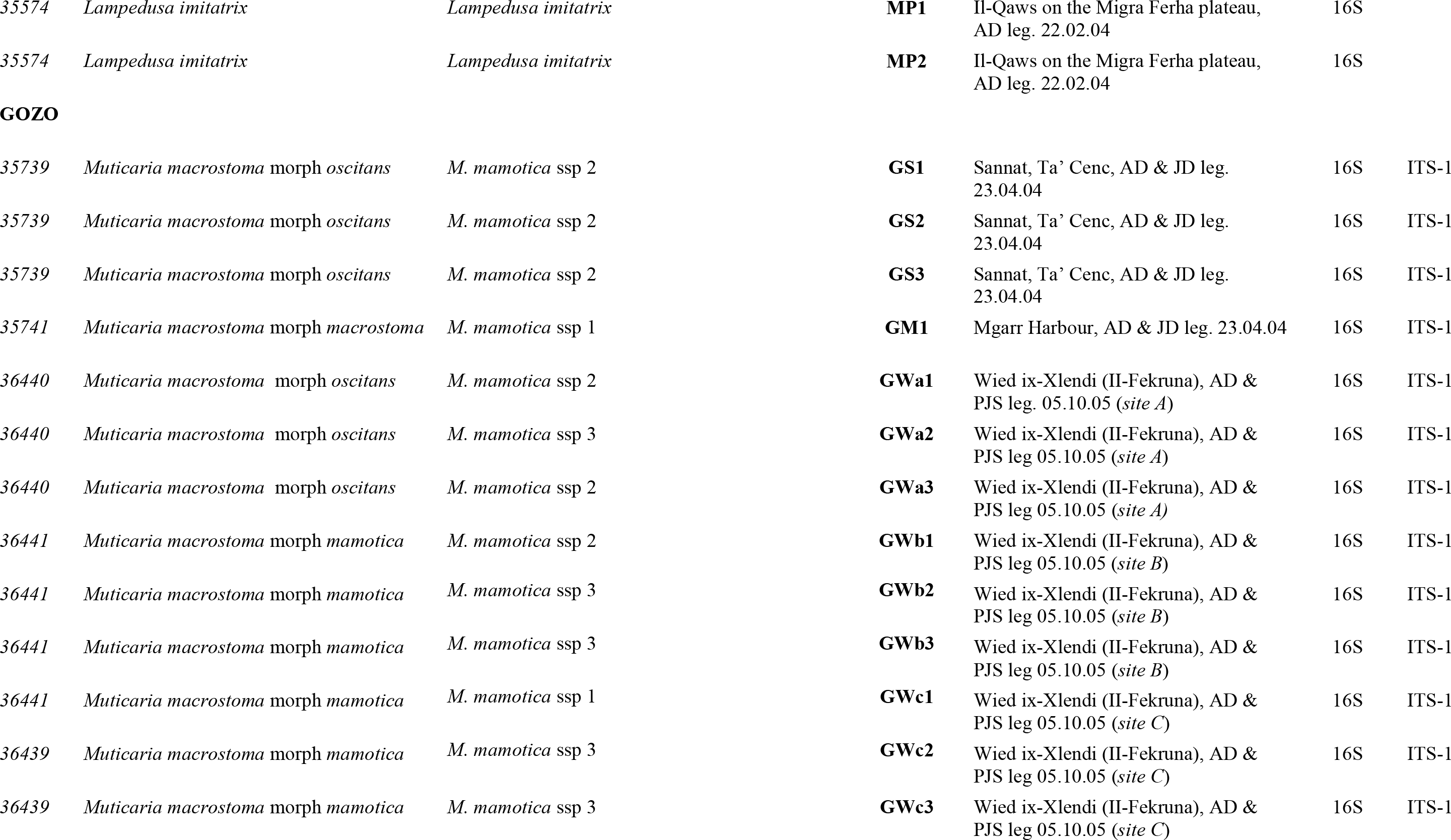
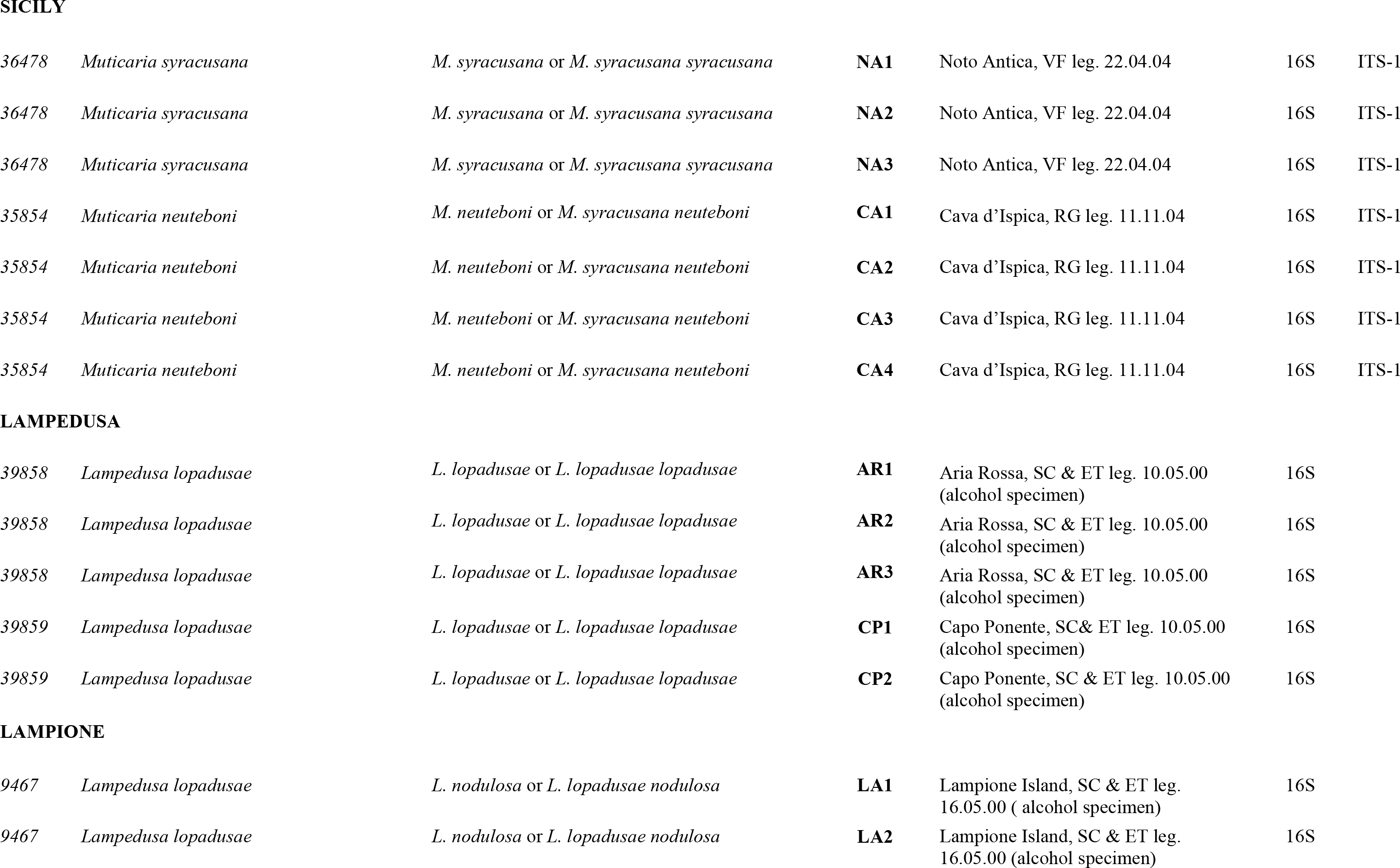
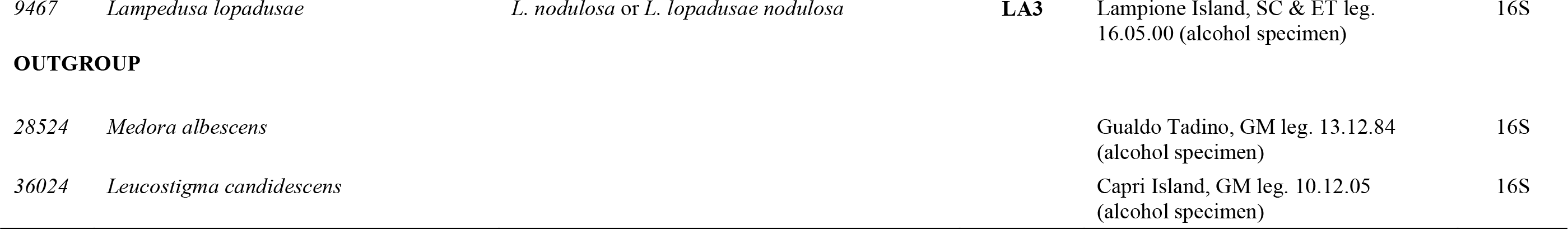
Material examined. FGC: F. Giusti collection inventory number, Department of Evolutive Biology, University of Siena. Collectors: AD Alan Deidun, ET Enrico Talenti, FG Folco Giusti, GM Giuseppe Manganelli, JD Joseph Debono, PJS Patrick J. Schembri, RG Rosario Grasso, SC Simone Cianfanelli, VF Viviana Fiorentino

| FGC | Current taxonomy<br>(Giusti et l., 1995) | revised taxonomy | Acronym | Locality, collectors and date | Gen bank<br>accession numbers |  |
| --- | --- | --- | --- | --- | --- | --- |
| MALTA |  |  |  |  |  |  |
| 35573 | *Muticaria macrostoma* morph *oscitans* | *M. oscitans* or *M. macrostoma oscitans* | MJ | Migra Ferha (car park), AD leg.<br>11.02.04 | 16S | ITS-1 |
| 35575 | *Muticaria macrostoma* morph *oscitans* | *M. oscitans* or *M. macrostoma oscitans* | MC1 | Clapham Junction, AD leg. 11.02.04 | 16S | ITS-1 |
| 35575 | *Muticaria macrostoma* morph *oscitans* | *M. oscitans* or *M. macrostoma oscitans* | MC2 | Clapham Junction, AD leg. 11.02.04 | 16S | ITS-1 |
| 35738 | *Muticaria macrostoma* morph *oscitans* | *M. oscitans* or *M. macrostoma oscitans* | MG1 | Ghar Lapsi (Wied Hoxt), AD & JD leg.<br>04.05.04 | 16S | ITS-1 |
| 35738 | *Muticaria macrostoma* morph *oscitans* | *M. oscitans* or *M. macrostoma oscitans* | MG2 | Ghar Lapsi (Wied Hoxt), AD & JD leg.<br>04.05.04 | 16S | ITS-1 |
| 35740 | *Muticaria macrostoma* morph *macrostoma* | *M. macrostoma* or *M. macrostoma macrostoma* | MMA1 | Mgarr (San Martin Valley), AD & JD<br>leg. 30.04.04 | 16S | ITS-1 |
| 35740 | *Muticaria macrostoma* morph *macrostoma* | *M. macrostoma* or *M. macrostoma macrostoma* | MMA2 | Mgarr (San Martin Valley), AD & JD<br>leg. 30.04.04 | 16S | ITS-1 |
| 35743 | *Muticaria macrostoma* morph *scalaris* | *M. scalaris* or *M. macrostoma scalaris* | MM1 | Mistra (Harrieq), AD & JD leg.<br>30.04.04 | 16S | ITS-1 |
| 35743 | *Muticaria macrostoma* morph *scalaris* | *M. scalaris* or *M. macrostoma scalaris* | MM2 | Mistra (Harrieq), AD & JD leg.<br>30.04.04 | 16S | ITS-1 |
| 33261 | *Lampedusa melitensis* | *Lampedusa melitensis* | LM | Rdum Il-Maddalena, FG, GM & PJS<br>leg. 27.11.87 (alcohol specimen) | 16S |  |
| 35574 | *Lampedusa imitatrix* | *Lampedusa imitatrix* | **MP1** | Il-Qaws on the Migra Ferha plateau,<br>AD leg. 22.02.04 | 16S |  |
| 35574 | *Lampedusa imitatrix* | *Lampedusa imitatrix* | **MP2** | Il-Qaws on the Migra Ferha plateau,<br>AD leg. 22.02.04 | 16S |  |
| **GOZO** |  |  |  |  |  |  |
| 35739 | *Muticaria macrostoma* morph *oscitans* | *M. mamotica* ssp 2 | **GS1** | Sannat, Ta' Cenc, AD & JD leg.<br>23.04.04 | 16S | ITS-1 |
| 35739 | *Muticaria macrostoma* morph *oscitans* | *M. mamotica* ssp 2 | **GS2** | Sannat, Ta' Cenc, AD & JD leg.<br>23.04.04 | 16S | ITS-1 |
| 35739 | *Muticaria macrostoma* morph *oscitans* | *M. mamotica* ssp 2 | **GS3** | Sannat, Ta' Cenc, AD & JD leg.<br>23.04.04 | 16S | ITS-1 |
| 35741 | *Muticaria macrostoma* morph *macrostoma* | *M. mamotica* ssp 1 | **GM1** | Mgarr Harbour, AD & JD leg. 23.04.04 | 16S | ITS-1 |
| 36440 | *Muticaria macrostoma* morph *oscitans* | *M. mamotica* ssp 2 | **GWa1** | Wied ix-Xlendi (II-Fekruna), AD &<br>PJS leg. 05.10.05 (*site A*) | 16S | ITS-1 |
| 36440 | *Muticaria macrostoma* morph *oscitans* | *M. mamotica* ssp 3 | **GWa2** | Wied ix-Xlendi (II-Fekruna), AD &<br>PJS leg 05.10.05 (*site A*) | 16S | ITS-1 |
| 36440 | *Muticaria macrostoma* morph *oscitans* | *M. mamotica* ssp 2 | **GWa3** | Wied ix-Xlendi (II-Fekruna), AD &<br>PJS leg 05.10.05 (*site A*) | 16S | ITS-1 |
| 36441 | *Muticaria macrostoma* morph *mamotica* | *M. mamotica* ssp 2 | **GWb1** | Wied ix-Xlendi (II-Fekruna), AD &<br>PJS leg 05.10.05 (*site B*) | 16S | ITS-1 |
| 36441 | *Muticaria macrostoma* morph *mamotica* | *M. mamotica* ssp 3 | **GWb2** | Wied ix-Xlendi (II-Fekruna), AD &<br>PJS leg 05.10.05 (*site B*) | 16S | ITS-1 |
| 36441 | *Muticaria macrostoma* morph *mamotica* | *M. mamotica* ssp 3 | **GWb3** | Wied ix-Xlendi (II-Fekruna), AD &<br>PJS leg 05.10.05 (*site B*) | 16S | ITS-1 |
| 36441 | *Muticaria macrostoma* morph *mamotica* | *M. mamotica* ssp 1 | **GWc1** | Wied ix-Xlendi (II-Fekruna), AD &<br>PJS leg 05.10.05 (*site C*) | 16S | ITS-1 |
| 36439 | *Muticaria macrostoma* morph *mamotica* | *M. mamotica* ssp 3 | **GWc2** | Wied ix-Xlendi (II-Fekruna), AD &<br>PJS leg 05.10.05 (*site C*) | 16S | ITS-1 |
| 36439 | *Muticaria macrostoma* morph *mamotica* | *M. mamotica* ssp 3 | **GWc3** | Wied ix-Xlendi (II-Fekruna), AD &<br>PJS leg 05.10.05 (*site C*) | 16S | ITS-1 |
| SICILY |  |  |  |  |  |  |
| 36478 | *Muticaria syracusana* | *M. syracusana* or *M. syracusana syracusana* | **NA1** | Noto Antica, VF leg. 22.04.04 | 16S | ITS-1 |
| 36478 | *Muticaria syracusana* | *M. syracusana* or *M. syracusana syracusana* | **NA2** | Noto Antica, VF leg. 22.04.04 | 16S | ITS-1 |
| 36478 | *Muticaria syracusana* | *M. syracusana* or *M. syracusana syracusana* | **NA3** | Noto Antica, VF leg. 22.04.04 | 16S | ITS-1 |
| 35854 | *Muticaria neuteboni* | *M. neuteboni* or *M. syracusana neuteboni* | **CA1** | Cava d'Ispica, RG leg. 11.11.04 | 16S | ITS-1 |
| 35854 | *Muticaria neuteboni* | *M. neuteboni* or *M. syracusana neuteboni* | **CA2** | Cava d'Ispica, RG leg. 11.11.04 | 16S | ITS-1 |
| 35854 | *Muticaria neuteboni* | *M. neuteboni* or *M. syracusana neuteboni* | **CA3** | Cava d'Ispica, RG leg. 11.11.04 | 16S | ITS-1 |
| 35854 | *Muticaria neuteboni* | *M. neuteboni* or *M. syracusana neuteboni* | **CA4** | Cava d'Ispica, RG leg. 11.11.04 | 16S | ITS-1 |

LAMPEDUSA
|  |  |  |  |  |  |
| --- | --- | --- | --- | --- | --- |
| 39858 | *Lampedusa lopadusae* | *L. lopadusae* or *L. lopadusae lopadusae* | **AR1** | Aria Rossa, SC & ET leg. 10.05.00<br>(alcohol specimen) | 16S |
| 39858 | *Lampedusa lopadusae* | *L. lopadusae* or *L. lopadusae lopadusae* | **AR2** | Aria Rossa, SC & ET leg. 10.05.00<br>(alcohol specimen) | 16S |
| 39858 | *Lampedusa lopadusae* | *L. lopadusae* or *L. lopadusae lopadusae* | **AR3** | Aria Rossa, SC & ET leg. 10.05.00<br>(alcohol specimen) | 16S |
| 39859 | *Lampedusa lopadusae* | *L. lopadusae* or *L. lopadusae lopadusae* | **CP1** | Capo Ponente, SC& ET leg. 10.05.00<br>(alcohol specimen) | 16S |
| 39859 | *Lampedusa lopadusae* | *L. lopadusae* or *L. lopadusae lopadusae* | **CP2** | Capo Ponente, SC & ET leg. 10.05.00<br>(alcohol specimen) | 16S |

LAMPIONE
|  |  |  |  |  |  |
| --- | --- | --- | --- | --- | --- |
| 9467 | *Lampedusa lopadusae* | *L. nodulosa* or *L. lopadusae nodulosa* | **LA1** | Lampione Island, SC & ET leg.<br>16.05.00 ( alcohol specimen) | 16S |
| 9467 | *Lampedusa lopadusae* | *L. nodulosa* or *L. lopadusae nodulosa* | **LA2** | Lampione Island, SC & ET leg.<br>16.05.00 (alcohol specimen) | 16S |

|  |  |  |  |  |  |
| --- | --- | --- | --- | --- | --- |
| 9467 | *Lampedusa lopadusae* | *L. nodulosa* or *L. lopadusae nodulosa* | **LA3** | Lampione Island, SC & ET leg.<br>16.05.00 (alcohol specimen) | 16S |

**OUTGROUP**
|  |  |  |  |  |  |
| --- | --- | --- | --- | --- | --- |
| 28524 | *Medora albescens* |  |  | Gualdo Tadino, GM leg. 13.12.84<br>(alcohol specimen) | 16S |
| 36024 | *Leucostigma candidescens* |  |  | Capri Island, GM leg. 10.12.05<br>(alcohol specimen) | 16S |

**Figure 2.**
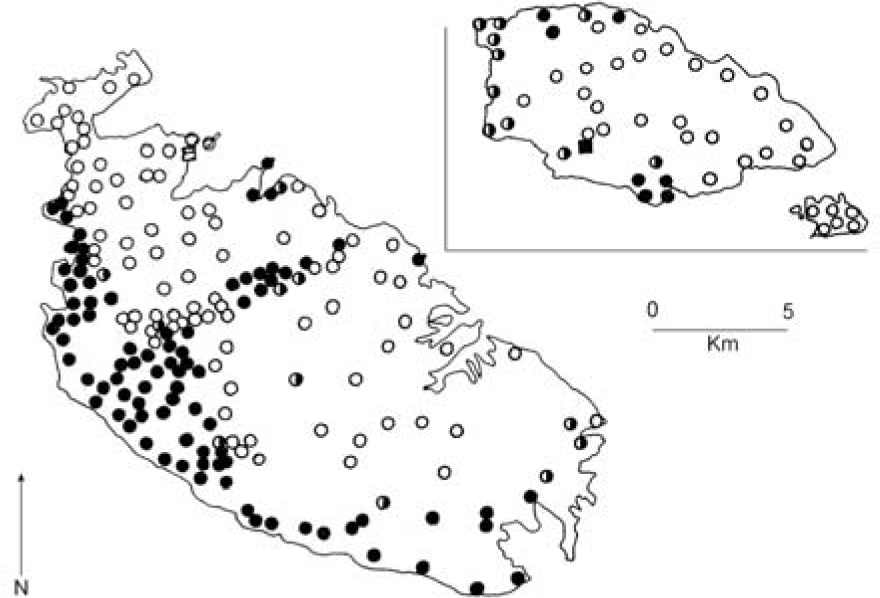
Distribution of *Muticaria* morphs in the Maltese Islands (modified from Thake, 1985). Filled square, *mamotica*; empty square *scalaris*; filled circles *oscitans*; empty circles *macrostoma*; half-filled circles, *macrostoma-oscitans* (population of difficult determination).

Interestingly, Thake (1985) and Holyoak (1986) reported an area of hybridization between *L. imitatrix* and *M. macrostoma*. Giusti (1995), re-examining the voucher specimens, demonstrated that some alleged hybrids belonged to *L. imitatrix* or *M. macrostoma*. However, some other specimens were real hybrids between species of *Lampedusa* and *Muticaria* (hybrids occur in a small area of a few tens of square metres, see Giusti et al., 1995 for details).

The Pelagian and Maltese archipelagos are located in the Sicilian Channel. Together with the eastern coast of Tunisia and the Hyblean region of Sicily, the Maltese and Pelagian islands are the only currently exposed parts of the Pelagian Block, which is the foreland margin of the African continental plate (Pedley, House & Waugh, 1978; Pedley 1990; Grasso & Pedley 1985; Gatt, 2007). The submerged parts of the Pelagian Block were exposed to form a land bridge or corridor (sensu Simpson, 1940) between northern Africa and Italy, via Sicily, starting ca 5.59 million years BP during the Messinian Salinity Crisis (Hsü, Ryan & Cita, 1973; Krijgsman et al, 1999, Krijgsman, 2002; CIESM, 2008), allowing biota from North Africa and the Italian peninsula to colonise the exposed land. Much of the Pelagian Block was gradually and intermittently submerged with the refilling of the Mediterranean at the end of the Miocene ca 5.33 million years BP (Krijgsman et al, 1999; Pedley et al., 2007) and it is generally held that Africa and Sicily have not been connected again since, although some evidence has now accumulated for an exchange of fauna between Africa and Sicily via a land bridge after the end of the Messinian Salinity Crisis (Stöck et al, 2008). Faunal exchange would almost certainly have occurred during the Plio-Pleistocene glaciations when the African palaeocoast would have repeatedly approached Sicily during sea level lowstands (Thiede, 1978; Rohling et al., 1998) facilitating jump dispersal across the narrow channel that separated the African and Sicilian palaeocoasts, a process that may have been assisted by any islands that were present in the Pleistocene Sicilian Channel acting as stepping stones for dispersal (Flemming et al., 2003); there are several banks in the present day Sicilian Channel that would have become exposed as islands during some of the Plio-Pleistocene marine regressions (British Admiralty, 2005, 2010).

The Maltese archipelago, located approximately 100 km from Sicily and 300 km from North Africa, consists of low islands aligned in a NW-SE direction. The three main islands of Malta (245.7 km^2^), Gozo (67.1 km^2^) and Comino (2.8 km^2^) are inhabited, and there are a number of uninhabited islets each less than 10 ha (Schembri, 1997). The islands are composed almost entirely of marine sedimentary rocks, mainly limestones of Oligo-Miocene age (30-5 million years BP) with some minor Quaternary deposits of terrestrial origin (Pedley, House & Waugh, 1976; Pedley, Hughes Clarke & Galea, 2002).

The Maltese Islands received their first influx of terrestrial biota during the Messinian when they were part of the Afro-Sicilian corridor. This biota was isolated following refilling of the Mediterranean and remained so during the Pliocene. However, further influxes of biota from Sicily occurred during the Pleistocene marine regressions either when the islands became connected to Hyblean Sicily by actual land bridges during the more extreme sea level lowstands, or by jump dispersal facilitated by a narrowing of the channel between Sicily and the Maltese Islands. With the present bathymetry, the drop in sea-level needed to connect Malta to Sicily is about 155 m but the maximum Pleistocene regression was of 120-130m (Bard, Hamelin & Fairbanks, 1990; Ferland, 1995; Rohling et al., 1999), however, tectonic uplift due to crustal rebound during regressions, and sedimentation (Gatt, 2007), may have caused the depth of the Sicilian-Maltese channel to vary in the past; on the other hand a drop of 100m in sea-level would narrow the present day Sicilian-Maltese channel to less than 14km wide (Hunt & Schembri, 1999).

In general, the Maltese biota resembles that of Sicily (Francini Corti & Lanza, 1973; Hunt & Schembri, 1999), although it also comprises a number of endemics (Giusti et al., 1995; Schembri, 2003). Only few species occur on Malta and North Africa and are not present in Sicily (Giusti et al., 1995; Schembri, 2003), although some of the Maltese endemic, or putatively endemic, species have a North African rather than a European affinity (Schembri, 2003).

The islands of Lampedusa and Linosa and the islet of Lampione constitute the Pelagian archipelago which lies on the northern edge of the African continental shelf. Lampedusa and Lampione consist of Oligo-Miocene limestones broadly similar to those of the Maltese group (Grasso & Pedley, 1985; Grasso, Pedley & Reuther, 1985), and like these islands, they arose in the Late Miocene (Torelli et al., 1995; André et al., 2002); Linosa has a volcanic origin. Like the Maltese Islands, the Pelagian archipelago was part of the exposed corridor of land between Africa and Europe during the Messinian, and remained isolated for a long time following the inundation of the Mediterranean at the end of the Messinian Salinity Crisis. However, during Quaternary marine lowstands it was connected to the African continent (Giraudi, 2004), but never to Sicily. The Pelagian biota is on the whole very disharmonic, and includes many endemics (Massa et al., 1995; Caruso, Noto La Diega & Bernini, 2005).

Numerous unanswered questions on the biogeography and taxonomy of the alopiine clausiliids of the Sicilian Channel, as well as the need for conserving the rarer members of this complex group of land snails, have prompted the present study. Our analysis includes representatives from nearly all taxa traditionally defined for this geographic area and constitutes a first attempt to build a phylogeny of the alopiinae radiation in the Sicilian Channel on the basis of both morphological and genetic data (shell morphology; sequencing of a fragment of the mitochondrial large ribosomal subunit, 16S rRNA, and the nuclear internal transcriber spacer 1, ITS-1 rRNA).

## Material and Methods

### SAMPLE COLLECTION

Information on sampling sites and specimens is summarized in Table 1 and Fig. 3.

All representatives of alopiine clausiliids reported from southeastern Sicily and the islands of the Sicily Channel were sampled. Since some taxa are endangered endemics, only a limited number of specimens was collected. Thus, whenever possible, alcohol preserved material was used.

**Figure 3.**
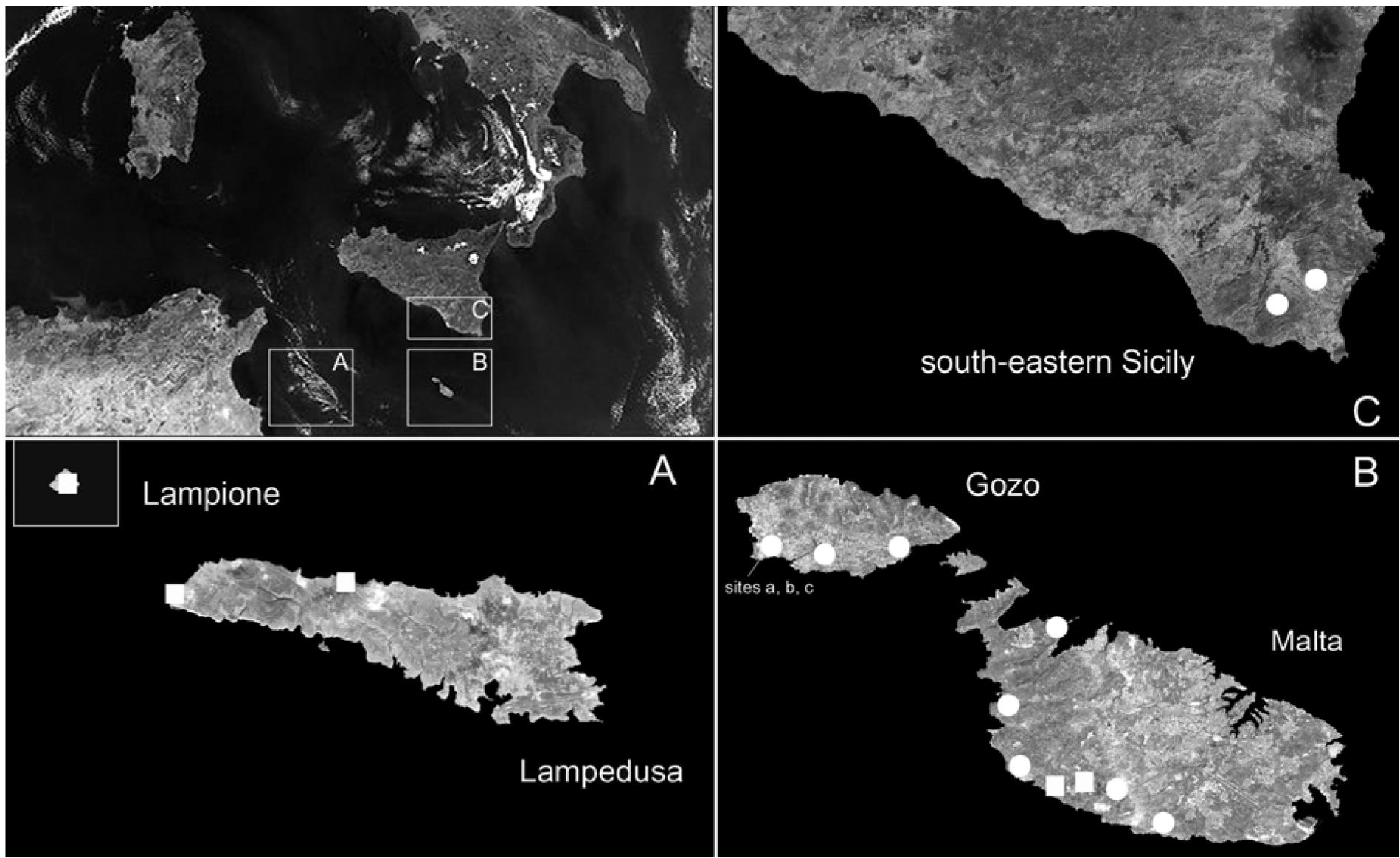
Sampling locations used in this study. See Table 1 for abbreviations.

Analyses were performed on operational groups determined on the basis of taxonomy and geographical distribution. Since genetic analyses (see below) separated robustly the *Lampedusa* specimens of Lampione islet from those of Lampedusa island, we considered this population separately, denoting it with the available name “*nodulosa*”. Consequently, the *Lampedusa* operational groups studied are: *imitatrix*, *lopadusae*, *melitensis* and *nodulosa*. Based on shell ribbing and shape, some populations of *Muticaria* from the Maltese Islands appeared to be intermediate between *macrostoma* and *oscitans* (central-south of Malta) or *mamotica* and *oscitans* (western Gozo). Thus we identified the following *Muticaria* operational groups: *macrostoma*, *macrostoma-oscitans*, *oscitans*, *mamotica*, *mamotica-oscitans*, *scalaris*, *syracusana* and *neuteboomi*.

Two different species were chosen as outgroups: *Medora italiana* (Küster, 1847) and *Leucostigma candidescens* (Rossmässler, 1835). Both species are generally thought to be closely related to the ingroup taxa, although their phylogenetic relationships have never been addressed.

### MORPHOLOGICAL DATA

Shell measurements were usually taken for ten specimens selected randomly from each locality, for a total of 56 specimens of *Lampedusa* and 470 of *Muticaria*. Only adult shells were used for measuring height (H), width (D) and the number of ribs (NR) on the penultimate whorl (Fig. 4). H and D were measured to the nearest 0.01 mm on shells positioned in apertural view, using an eyepiece micrometer fitted in a light microscope (Wild M5A) (Fig. 4).

**Figure 4.**
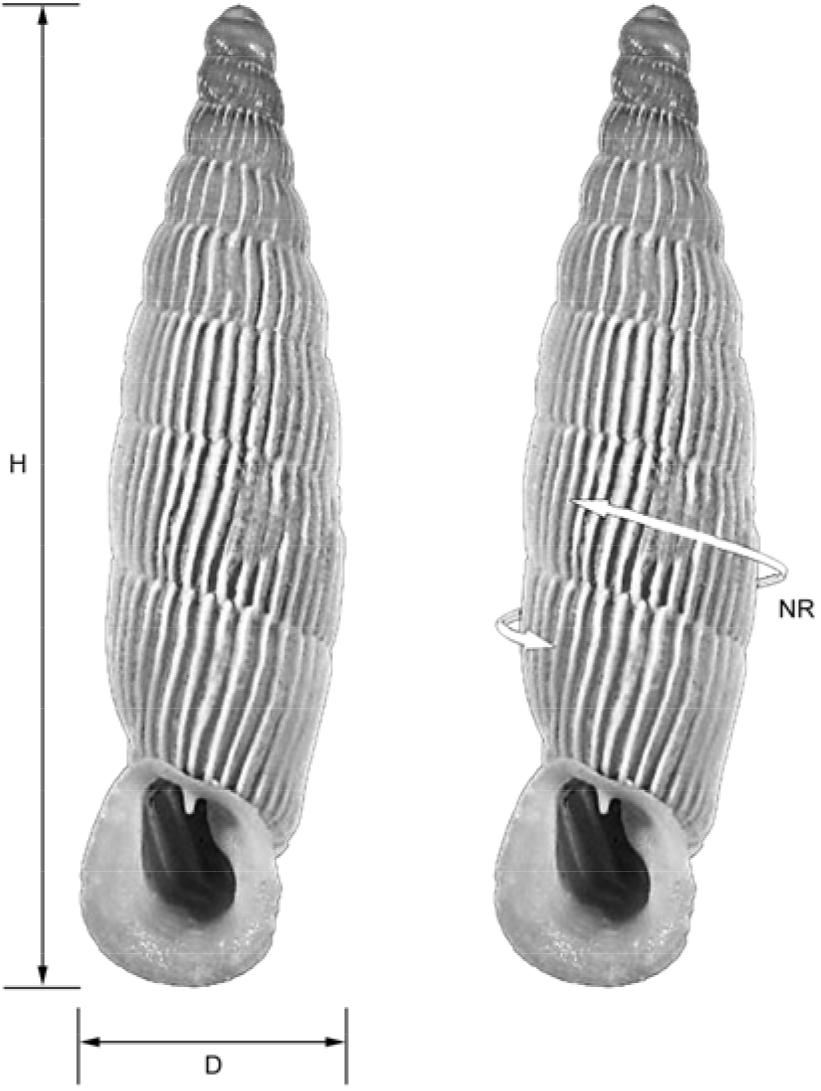
Shell shapes and measurements. Abbreviations: D, diameter; H, height.

Genital measurements were recorded for five sexually developed specimens randomly selected from each locality, for a total of 25 specimens of *Lampedusa* and 40 of *Muticaria*. Specimens were dissected under a light microscope (Wild M5A) using fine-pointed watchmaker’s tweezers. Eight linear variables (Fig. 5) were measured on isolated genitalia using an eyepiece micrometer fitted in a light microscope (accurate to 0.01 mm).

**Figure 5.**
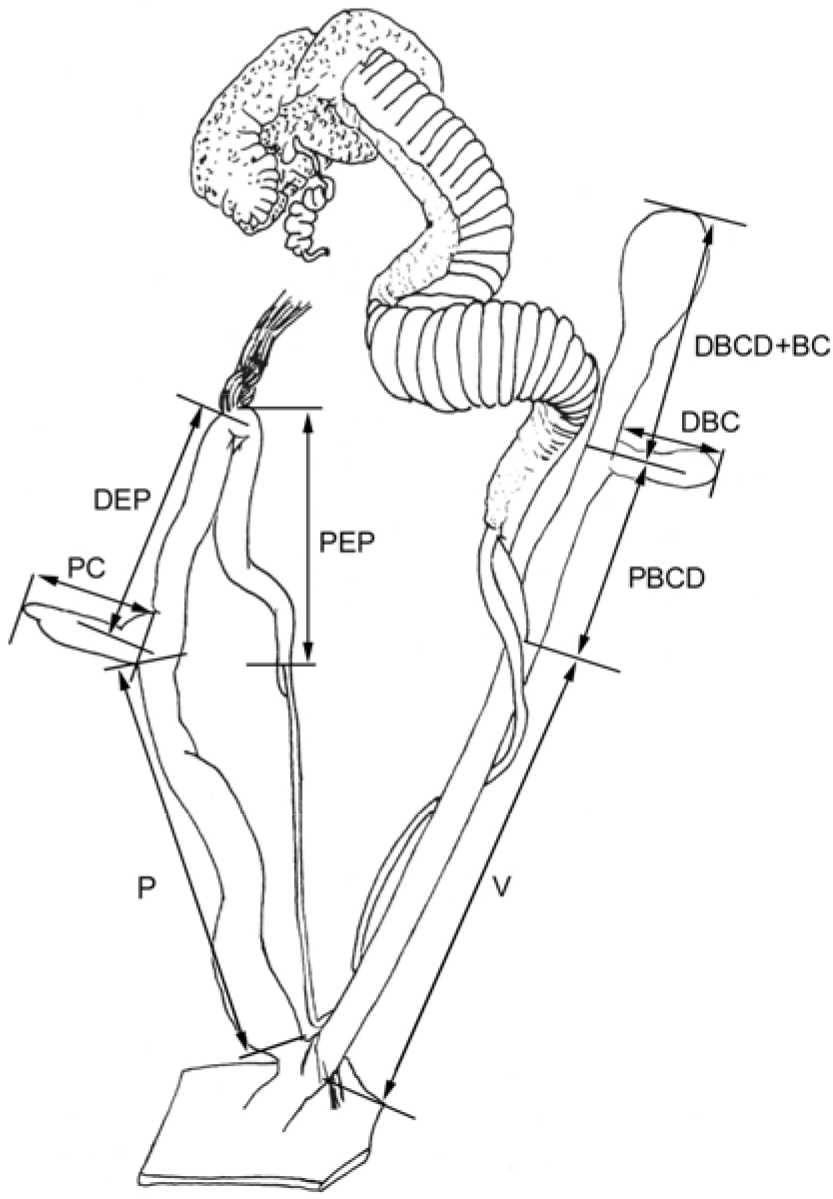
Distal genitalia of *Muticaria*: outline and measurements. Abbreviations: BC bursa copulatrix, DBC diverticulum of bursa copulatrix duct, DEP distal epiphallus, P penis, PBCD proximal bursa copulatrix duct, PC penial caecum, PEP proximal epiphallus, V vagina.

### ANALYSIS OF MORPHOLOGICAL DATA

Morphological variables were log-transformed to obtain linear relationships, when necessary.

Two-way analysis of variance (ANOVA) was performed on shell measurements and number of ribs. The *a posteriori* Tukey test (α = 0.05) was used to check group significance. All the analyses were run for the *Muticaria* data set, for the *Lampedusa* data set, and for the two data sets combined.

Discriminant Function Analysis (DFA) was then performed considering all measured genital variables. The analysis was run with groups defined *a priori* (“operational groups” and “islands”). With this analysis we assessed which measurements contributed to discrimination of groups defined *a priori*. The sequential chi-square test was used to quantify the extent to which each discriminant function significantly separated groups and structure, and canonical coefficient tables were used to establish the contribution of each measurement to the first two discriminant functions.

All calculations were made using R-package version 2.3.0 (R Development Core Team, 2006). The STATISTICA 5 (StatSoft Inc., Tusla, USA) package was used to run the DFA analyses.

### DNA EXTRACTION, PCR AND SEQUENCING

A total of 11 specimens of *Lampedusa* from five sites and 30 of *Muticaria* from 12 sites were studied. Specimens of *Lampedusa* were representatives of all taxa (except *L. imitatrix gattoi* from Filfla islet); specimens of *Muticaria* were representatives of all operational groups.

Total genomic DNA was extracted from foot muscle of fresh or alcohol preserved specimens using the C-TAB buffer (0.1M Tris-HCl pH 8.0, 1.4 M NaCl, 0.02 M EDTA, 2% CTAB, 0.2% 2-mercaptoethanol) and subsequent standard phenol-chloroform/ethanol extraction (Hillis et al., 1996).

For all the sampled snails, a fragment of the mitochondrial gene encoding for the large ribosomal subunit (16S rDNA) was polymerase chain reaction (PCR) amplified using the primer pair 5’-CGATTTGAACTCAGATCA-3’ (Simon et al., 1994) and 5’-GTGCAAAGGTAGCATAATCA-3’(Gantenbein et al., 1999). In addition, the nuclear ribosomal gene cluster encompassing the ribosomal internal transcribed spacer (ITS-1) was sequenced in specimens of the genus *Muticaria*, using primers annealing to flanking regions of the 18S and the 5.8S (CS249, 5’-TCGTAACAAGGTTTCCG-3’ and DT421, 5’-GCTGCGTTCTTCATCG-3’; Schlötterer et al., 1994).

All PCR reactions were carried out in a total volume of 50 μl under the following conditions: 95°C for 20’’, 55°−52°C for 30’’ and 72°C for 30’’ (repeated for 25 cycles), plus a final extension step at 72°C for 5’. Reaction products were isolated on 1% agarose gel, excised under long-wavelength UV light, and purified using a “Nucleospin extract” (Genenco™) column kit. Both strands of the amplified fragments were directly cycle-sequenced using the same amplificationprimers and the CEQ dye terminator cycle sequencing kit. DNA sequences were then electrophoresed on a CEQ 8000XL (Beckman Coulter™). DNA sequences have been deposited in the GenBank database (see Table 1 for GenBank references - *will be submitted on acceptance*).

### ANALYSIS OF DNA SEQUENCES

The 16S rDNA sequences were aligned and checked with Clustal X (version 1.8, Thompson et al. 1997) and easily aligned by eye where necessary. Phylogenetic relationships were conducted on the 16S mitochondrial dataset using Maximum Parsimony and Bayesian inference. Parsimony analyses were performed with PAUP* (version 4.0, Swofford, 2001) using the heuristic search option with equal weighting of all characters (ACCTRAN character-state optimisation, 100 random stepwise additions, TBR branch-swapping algorithm) (Farris, 1970). To assess the robustness of the phylogenetic hypotheses, 1000 bootstrap replicates were performed (Felsenstein, 1985).

Prior to Bayesian analysis, we determined an appropriate model of sequence evolution using MRMODELTEST (vers. 2.2, Nylander, 2004). Bayesian analysis was then carried out with MRBAYES (version 3.1, Huelsenbeck & Ronquist, 2001; Ronquist & Huelsenbeck, 2003) using the same model as estimated from MRMODELTEST. MRBAYES was run for 2 million generations with a sampling frequency of 100 generations and one cold and three heated Markov chains.

Reached stationarity was evaluated by plotting the likelihood scores of sampled trees against generation time. Trees generated before the stationarity phase were discarded as “burn-in” and posterior probability values for each node were calculated based on the remaining sampled trees.

The nuclear ITS-1 region was easily aligned by eye using the program BIOEDIT (version 7.0, Hall 1999). Relationships within congeneric populations were then inferred by constructing a median-joining network (Bandelt, Forster & Röhl, 1999) using the program NETWORK 4.1.0 (www.fluxus-engineering.com, Fluxus Technology Ltd.).

## Results

### MORPHOLOGICAL ANALYSIS

ANOVA performed on H, D, D/H and NR of the shells of *Lampedusa* and *Muticaria* revealed significant differences between groups defined *a priori* (Tables 2–5; Figs. 6–7).

**Figure 6.**
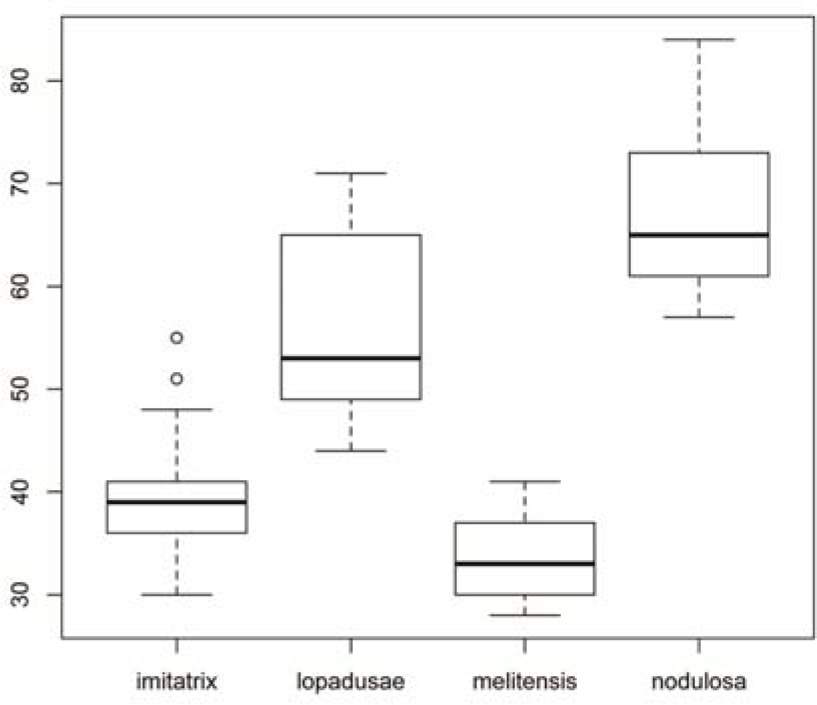
Box plot of the number of ribs in *Lampedusa*.

**Figure 7.**
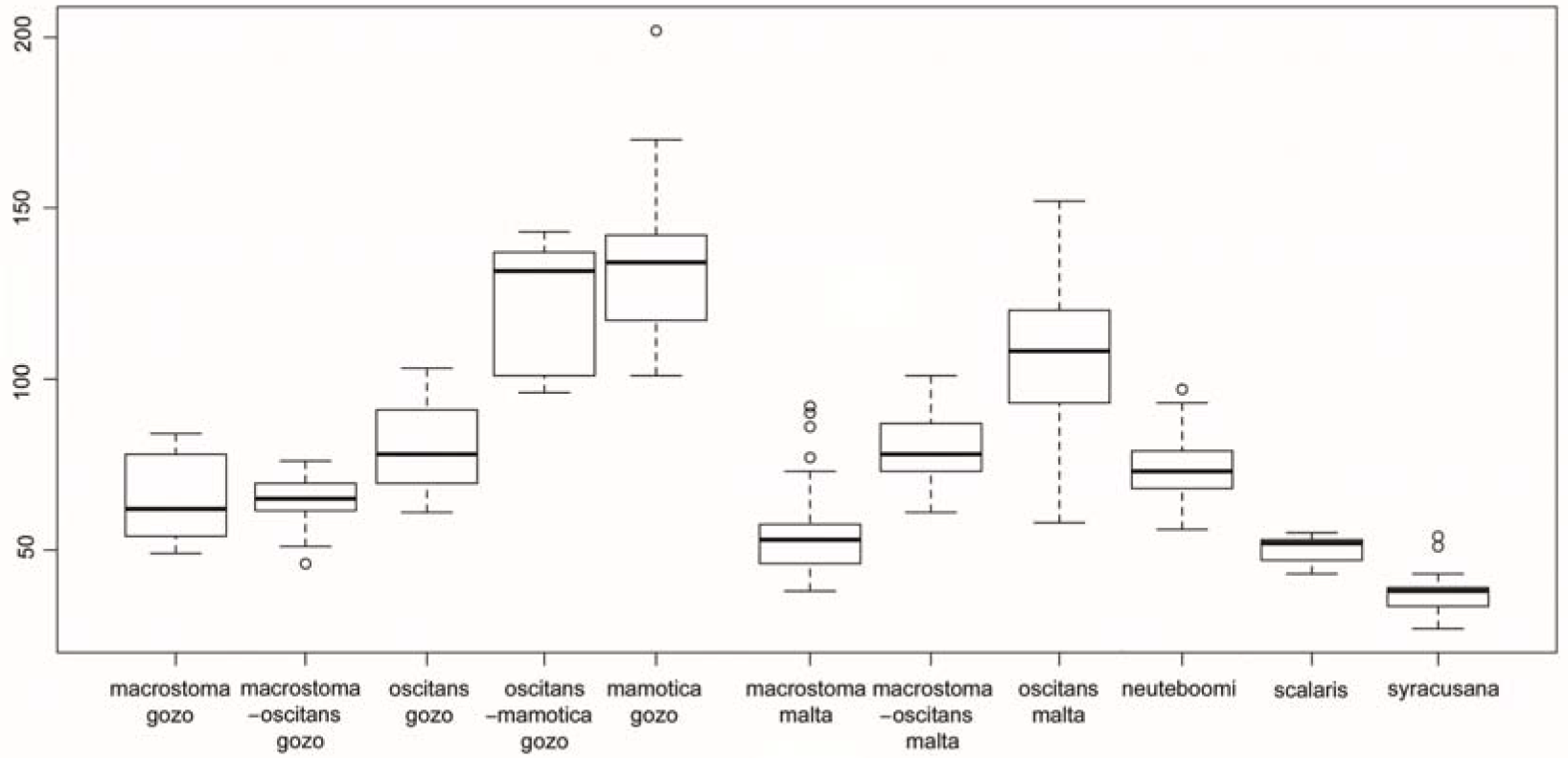
Box plot of the number of ribs in *Muticaria*.

**Table 2.** Two-way analysis of variance (ANOVA) on numbers of ribs in shell of *Lampedusa*.

| Taxa | diff | lwr | upr | p |
| --- | --- | --- | --- | --- |
| imitatrix-melitensis | 6.51 | -0.22 | 13.25 | 0.06 |
| lopadusae-melitensis | 22.3 | 14.20 | 30.39 | 0 |
| nodulosa-melitensis | 33.5 | 25.40 | 41.59 | 0 |
| lopadusae-imitatrix | 15.78 | 9.05 | 22.52 | 0 |
| nodulosa-imitatrix | 26.98 | 20.25 | 33.72 | 0 |
| nodulosa-lopadusae | 11.2 | 3.10 | 19.29 | 0 |

**Table 3.** Two-way analysis of variance (ANOVA) on the ratio D/H in *Lampedusa*.

| Taxa | diff | lwr | upr | p |
| --- | --- | --- | --- | --- |
| lopadusae-melitensis | 0.02847848 | -0.0496073 | 0.10656422 | 0.77 |
| nodulosa-melitensis | 0.03193357 | -0.0461522 | 0.11001931 | 0.70 |
| imitatrix-melitensis | 0.03237076 | -0.0326005 | 0.09734203 | 0.55 |
| nodulosa-lopadusae | 0.00345509 | -0.0746307 | 0.08154083 | 0.99 |
| imitatrix-lopadusae | 0.00389228 | -0.061079 | 0.06886355 | 0.99 |
| imitatrix-nodulosa | 0.00043719 | -0.0645341 | 0.06540845 | 0.99 |

**Table 4.**
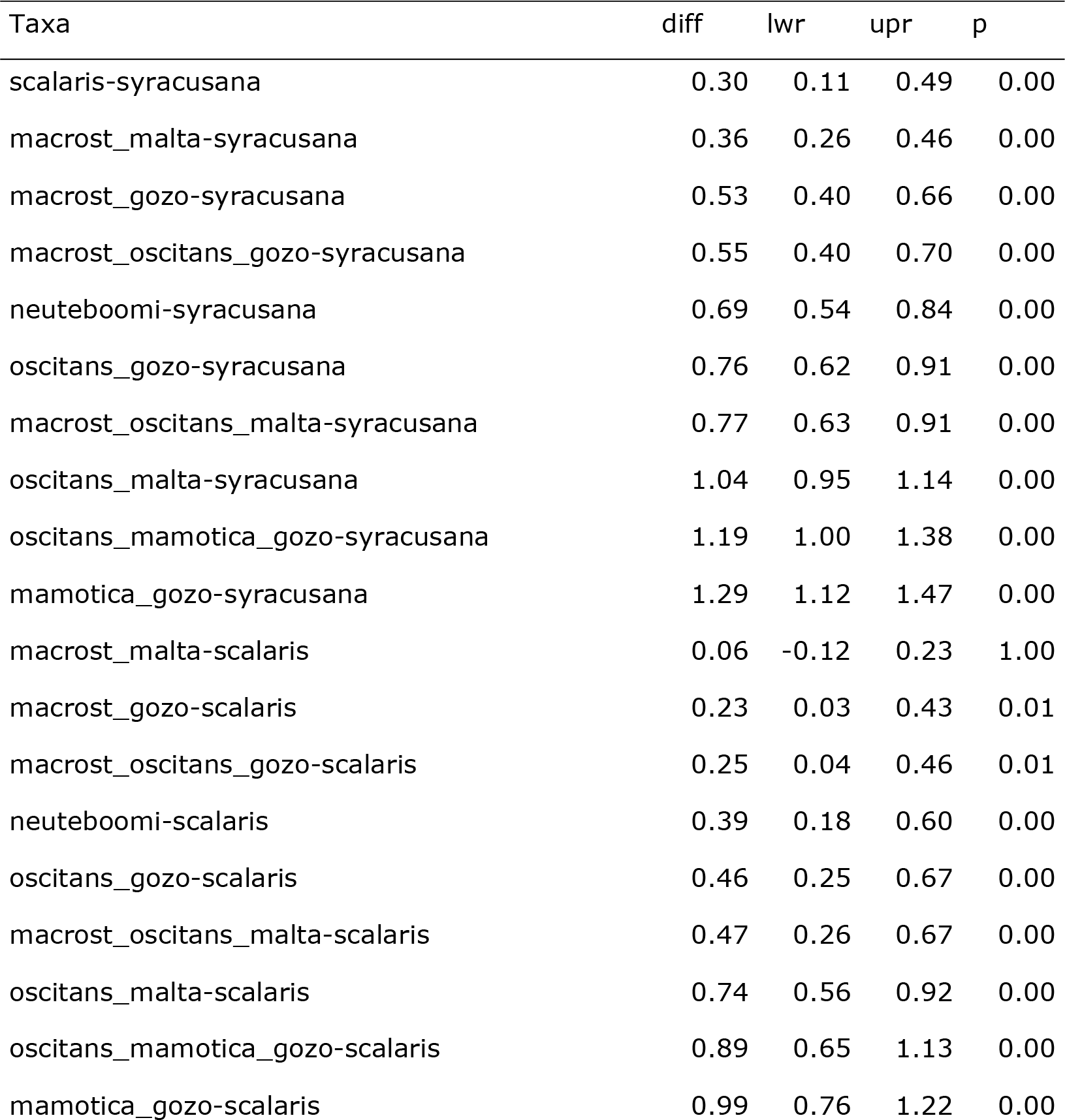
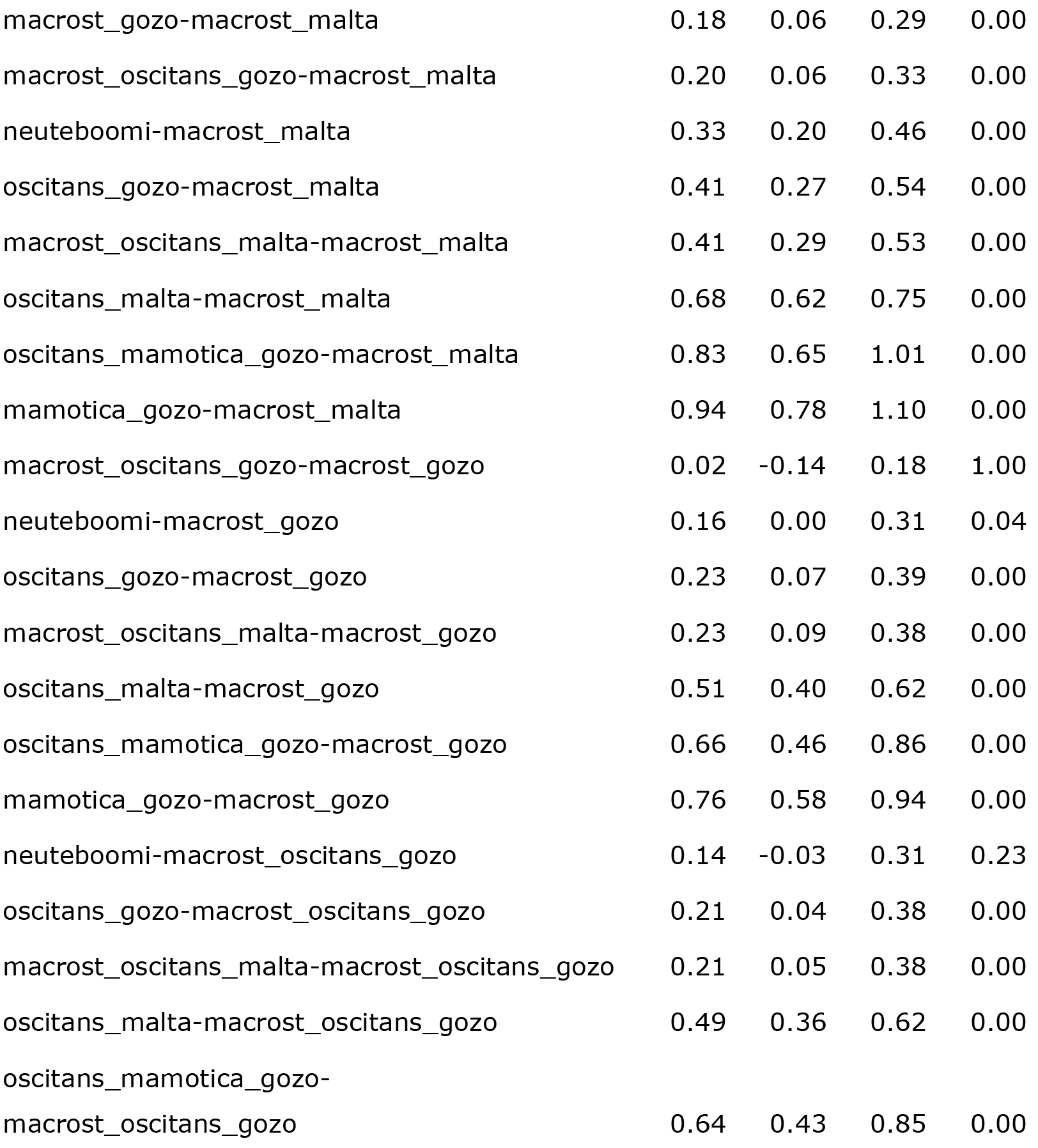
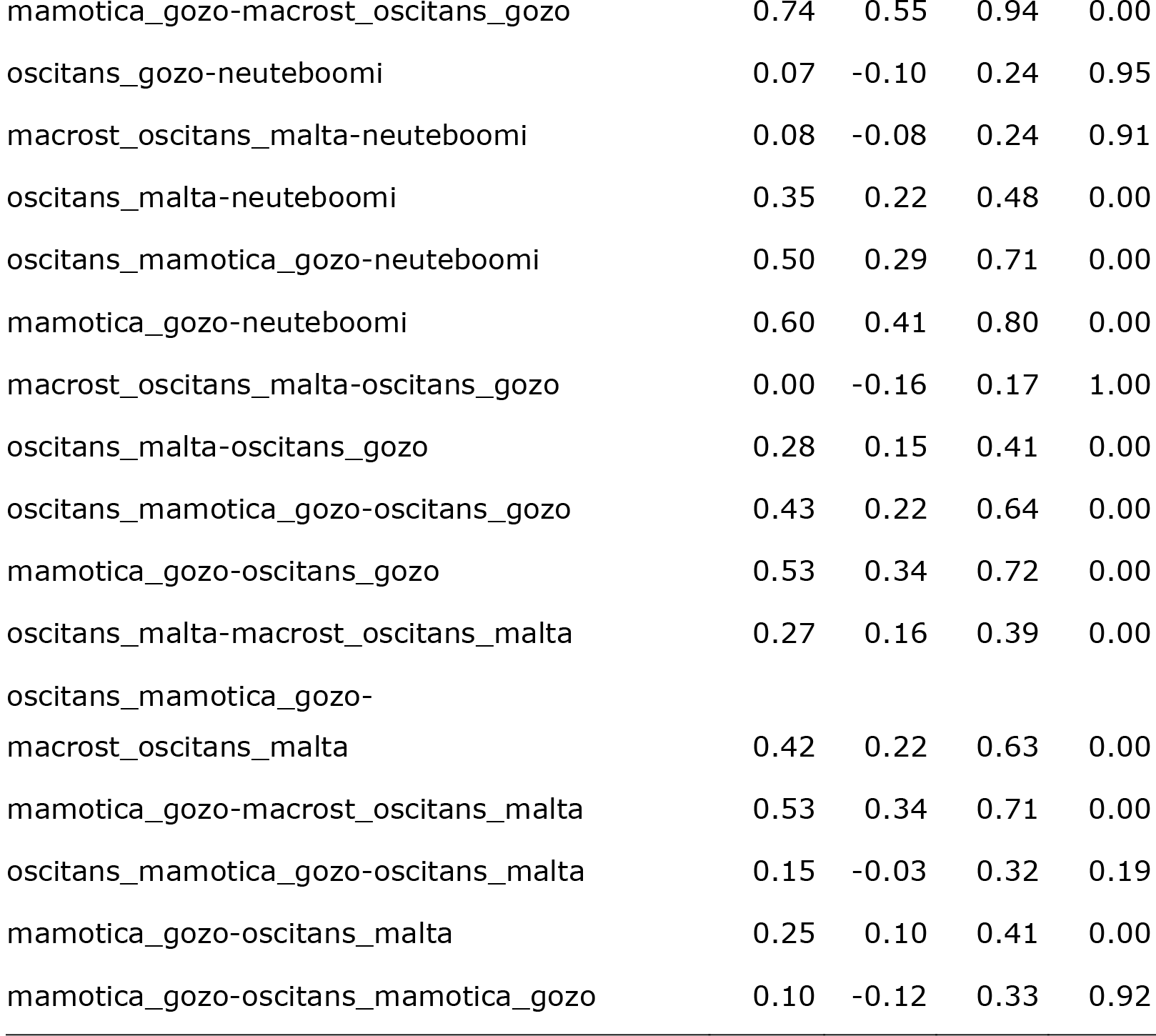
Two-way analysis of variance (ANOVA) on numbers of ribs in shell of Muticaria.

**Table 5.**
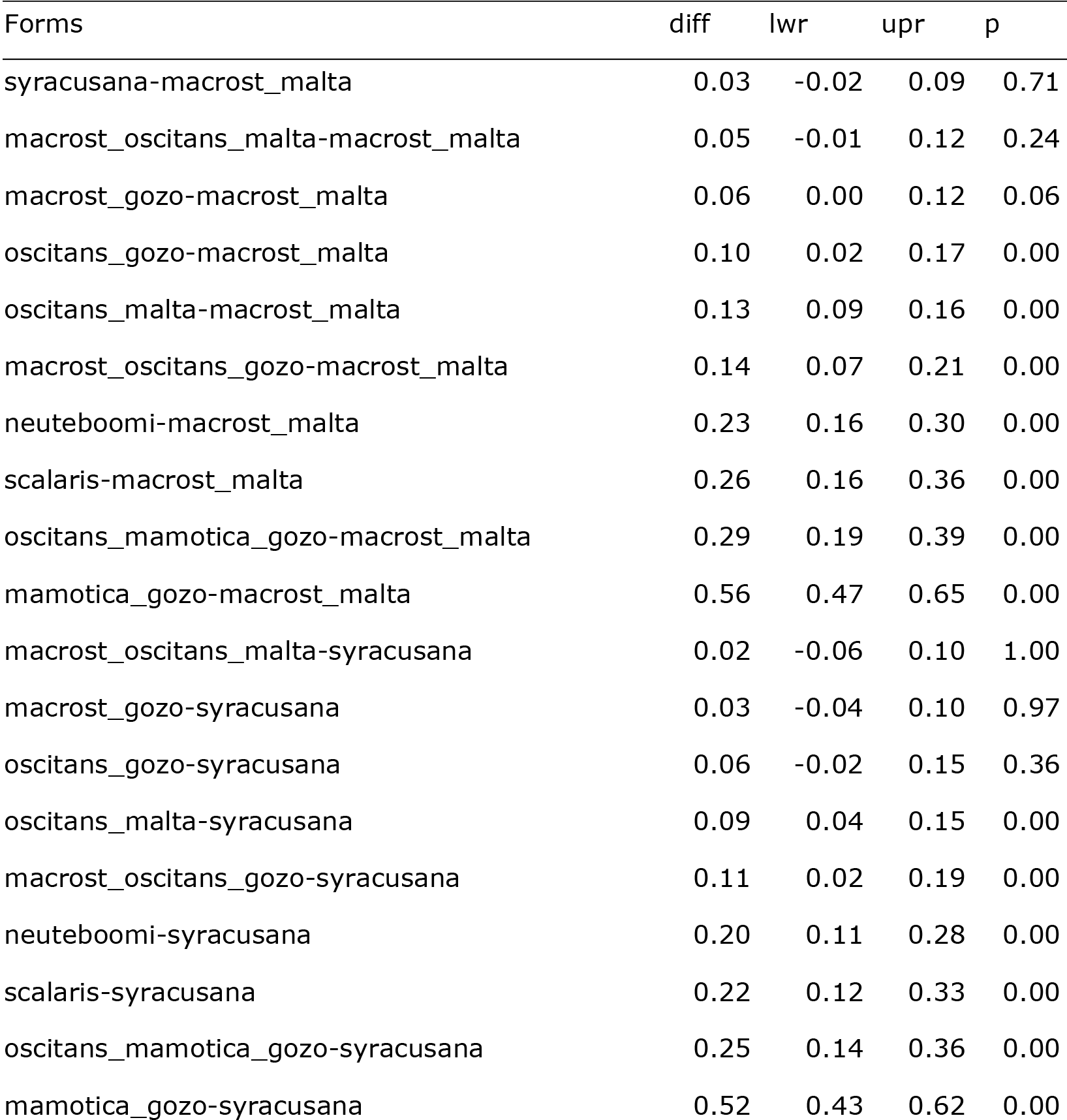
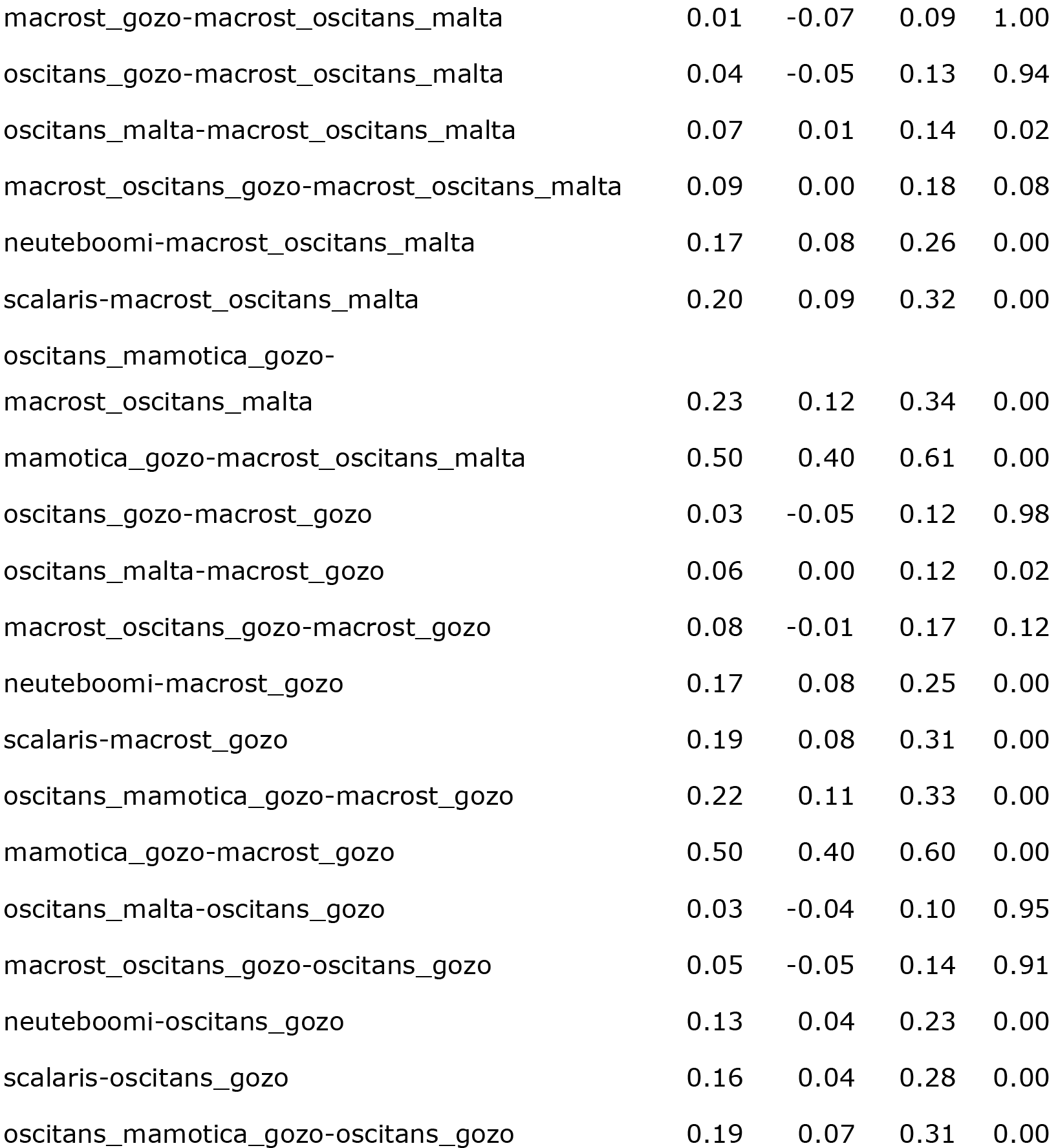
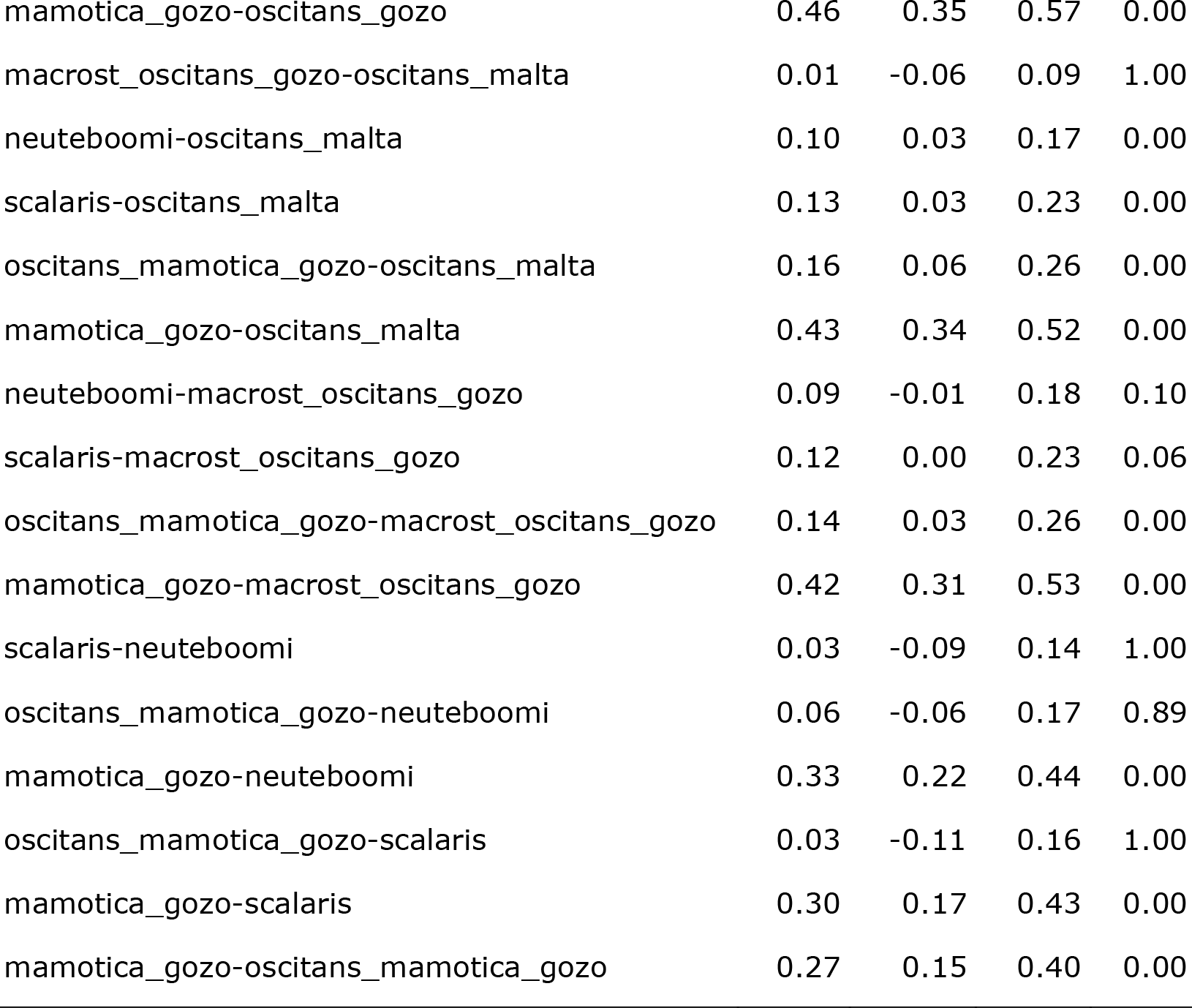
Two-way analysis of variance (ANOVA) on the ratio D/H in *Muticaria*.

ANOVA revealed that NR clearly distinguished the *Lampedusa* of Lampedusa from the *Lampedusa* of Malta and distinguished the *Lampedusa* of Lampione from all the other *Lampedusa*.

ANOVA showed that NR in *Muticaria* was also reliable for recognising the forms from each island, excluding the following groups: (1) Maltese *macrostoma* and *scalaris*; (2) Maltese *macrostoma-oscitans* and *neuteboomi*; (3) Maltese *macrostoma-oscitans* and Gozitan *oscitans*; (4) Gozitan *oscitans* and *neuteboomi*; (5) Gozitan *macrostoma-oscitans* and Gozitan *macrostoma*; (6) Gozitan *macrostoma-oscitans* and *neuteboomi*; (7) *oscitans-mamotica* and *mamotica*; (8) *oscitans-mamotica* from Gozo and Maltese *oscitans*.

DFA on the genital variables of *Lampedusa* revealed that the first discriminant function accounted for 97% of the variance while the second accounted only for 3%. The highest loadings on the first function were DBC (-6.33) and PEP (12.26). ANOVA on DBC and PEP showed significant differences between species. *L. imitatrix* was significantly different from *L. melitensis* and *L. lopadusae* for DBC; while *L. lopadusae* was significantly different from *L. imitatrix* and *L. melitensis* for PEP. Analyses on ratios of all the variables considered did not result in significant differences among species.

DFA on genital variables of *Muticaria* showed that the first discriminant function accounted for 69% of the variance and the second accounted for 28%. The highest loadings on the first function were: PC (- 1.02), DBC (-0.73), P (0.77) and PEP (0.57). ANOVA on these variables did not clearly distinguish species but only some pairs of taxa. In particular, F distinguished *syracusana* and *scalaris*, and P distinguished *syracusana* and *oscitans*.

### SEQUENCE CHARACTERISTICS

A total of 413 bp of the mitochondrial 16S rRNA was sequenced. Few indels were found in the alignment. However, removal or inclusion of indels in the phylogenetic analyses (indels were counted as one single mutation each, regardless of size) did not result in significant differences in tree topologies. There were a total of 152 variable characters, 96 of which were parsimony informative.

Base composition was homogeneous (*X*^*2*^ = 29.1, df = 120, P = 1.0), but skewed toward a deficiency in guanine (16.9%) and cytosine (13.5%), as expected for mitochondrial genes (Simon et al., 1994).

The nuclear dataset was limited to the subset including representatives of *Muticaria*. Alignment of the 30 ITS-1 nuclear sequences resulted in a matrix with 495 nucleotide positions (including gaps), providing 13 different nuclear variants. On the agarose gels, no evidence of intra-individual length variation was observed. There were 14 variable sites, 9 parsimony informative sites and 11 possible insertions or deletions. The distribution of indels seemed to be diagnostic for specific ITS-1 geographic variants, allowing, in particular, distinction between specimens from Sicily and those from Malta and Gozo. Nucleotide composition was skewed towards an increase of G+C (total = 59.1%).

### PHYLOGENETIC RELATIONSHIPS

Parsimony analysis from 16S rDNA sequence data produced three equally parsimonious trees (tree length=330, CI= 0.660, RI= 0.861) showing essentially the same topology.

The most appropriate model selected by MRMODELTEST was HKY+Г+I (Hasegawa, Kishino & Yano, 1985). Plots of the -ln likelihood scores over generation time showed that stable parameter estimates were obtained after approximately 300 trees (=30.000 generations). Therefore, only trees sampled after this burn-in period were used to determine posterior probabilities of model parameters (bpp), branch lengths and clades and to generate a 50% majority-rule consensus tree with PAUP*. The analysis was repeated several times with the same settings and it always generated similar results.

Parsimony and Bayesian analyses produced largely congruent results (Fig. 8), but with some topological differences (Fig. 9). Overall, both methods separated the *Lampedusa* and *Muticaria* haplotypes into two well distinct and supported lineages.

**Figure 8.**
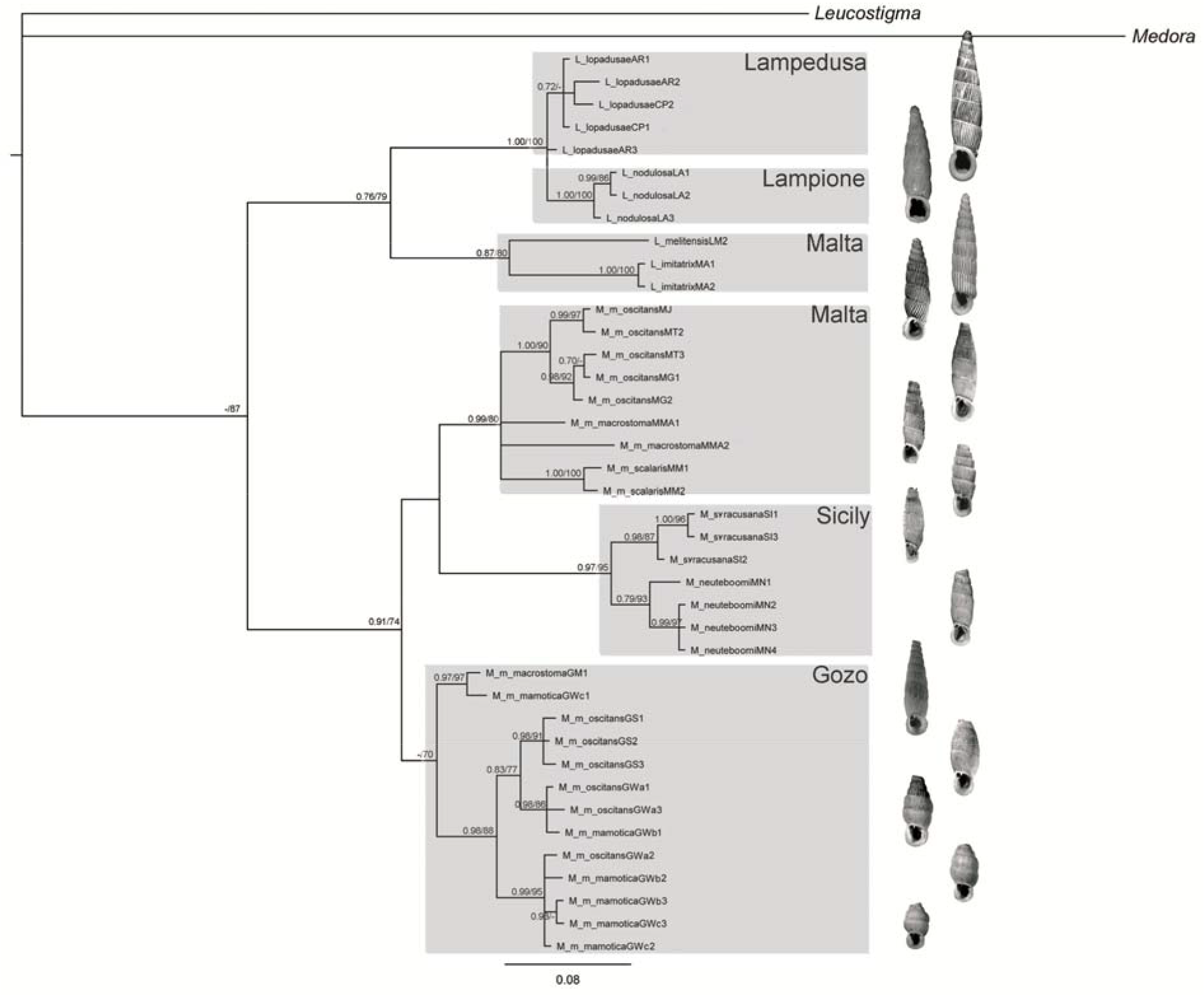
Consensus tree (50% majority rules) from Bayesian analysis based on mitochondrial 16S sequence data. Numbers at nodes represent MP bootstrap values >70% and posterior clades probability of each clade.

Populations grouped in the *Lampedusa* clade were further subdivided into two supported assemblages corresponding to geographic areas. The first subclade grouped specimens from Malta in two distinct lineages, corresponding to *L. melitensis* and *L. imitatrix*. The second subclade grouped all specimens from Lampedusa (*L. lopadusae* / *L. l. lopadusae*) and Lampione islet (*L. nodulosa* / *L. lopadusae nodulosa*). Remarkably, the three individuals from Lampione islet were robustly distinguished from those of Lampedusa island.

Within representatives of *Muticaria*, both parsimony and Bayesian reconstructions clearly defined three main groups. The first included all specimens from Sicily, resolved into two subclades, one for *M. neuteboomi* (site CI) and the other for *M. syracusana* (site NA). A second lineage included all individuals from Malta. Within this group, all specimens corresponding to *oscitans* (sites MJ, MG and MG) were grouped in a distinct and well supported monophyletic lineage. Specimens of *scalaris* (site MM) were also resolved as a separate monophyletic subgroup, while individulas of *macrostoma* (site MA) were in two distinct lineages characterized by distantly related haplotypes. The third main assemblage included all the specimens from Gozo. The *macrostoma* specimens GM1 and the *mamotica* specimens GWc1 formed a supported and distinct clade. According to the Bayesian analysis, these latter haplotypes (GM1 and GWc1) were unresolved within the *Muticaria* assemblage. All the other specimens from Gozo were grouped in a clade subdivided into two subclades, containing *oscitans* and *mamotica* haplotypes.

Relationships between the three main *Muticaria* groups were differently defined by the Baysian and Maximum Parsimony analyses (Fig. 9). Bayesian reconstructions placed haplotypesfrom Gozo as a sister group of the Sicilian + Maltese lineages, but with a scarce support (68%). On the contrary, Parsimony analysis suggested the Sicilian clade as sister group of the Gozo + Malta lineage (81%).

**Figure 9.**
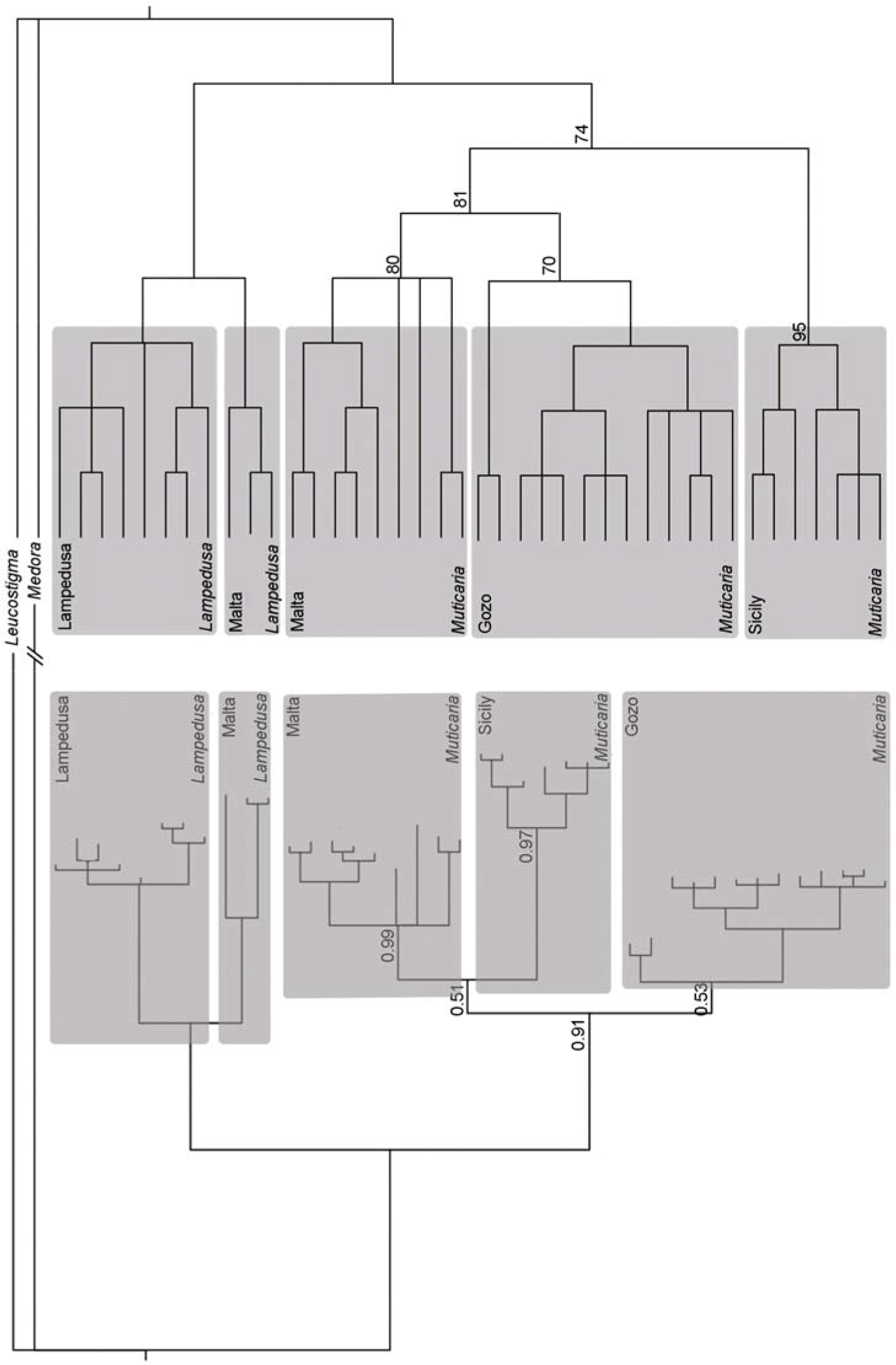
Comparison between consensus trees (50% majority rules) from Bayesian analysis and Maximum Parsimony analysis based on mitochondrial 16S sequence data. Numbers at basal nodes of *Muticaria* clades represent posterior clade probability and MP bootstrap values.

The median-joining network from the nuclear data set did not robustly distinguish the *Muticaria* sequences as did mtDNA data, but recognised three major groups: Sicily (sites CI and NA), south-east Malta (MJ, MG and MC) and north-west Malta + Gozo (Fig. 10). Populations from Sicily (CA and NA) showed two distinct nuclear variants close to each other (three mutational steps) and connected to those from southeastern Malta (9 mutational steps). The network was also indicative of a clear split between populations from southern-central-western Malta (MJ, MG and MC) and those from northwestern Malta (MMA and MM) + Gozo (GW, GS and GM). Relationships within this latter group were not well resolved. The two specimens of *macrostoma* from Malta (site MMA) shared the same sequence with an individual from Gozo (GWb2) and were separated by one mutational step from *scalaris* from Malta (site MM). The remaining specimens from Gozo were close to nuclear variants from Malta and showed an overall star-like pattern with two most common nuclear variants shared among different local populations.

**Figure 10.**
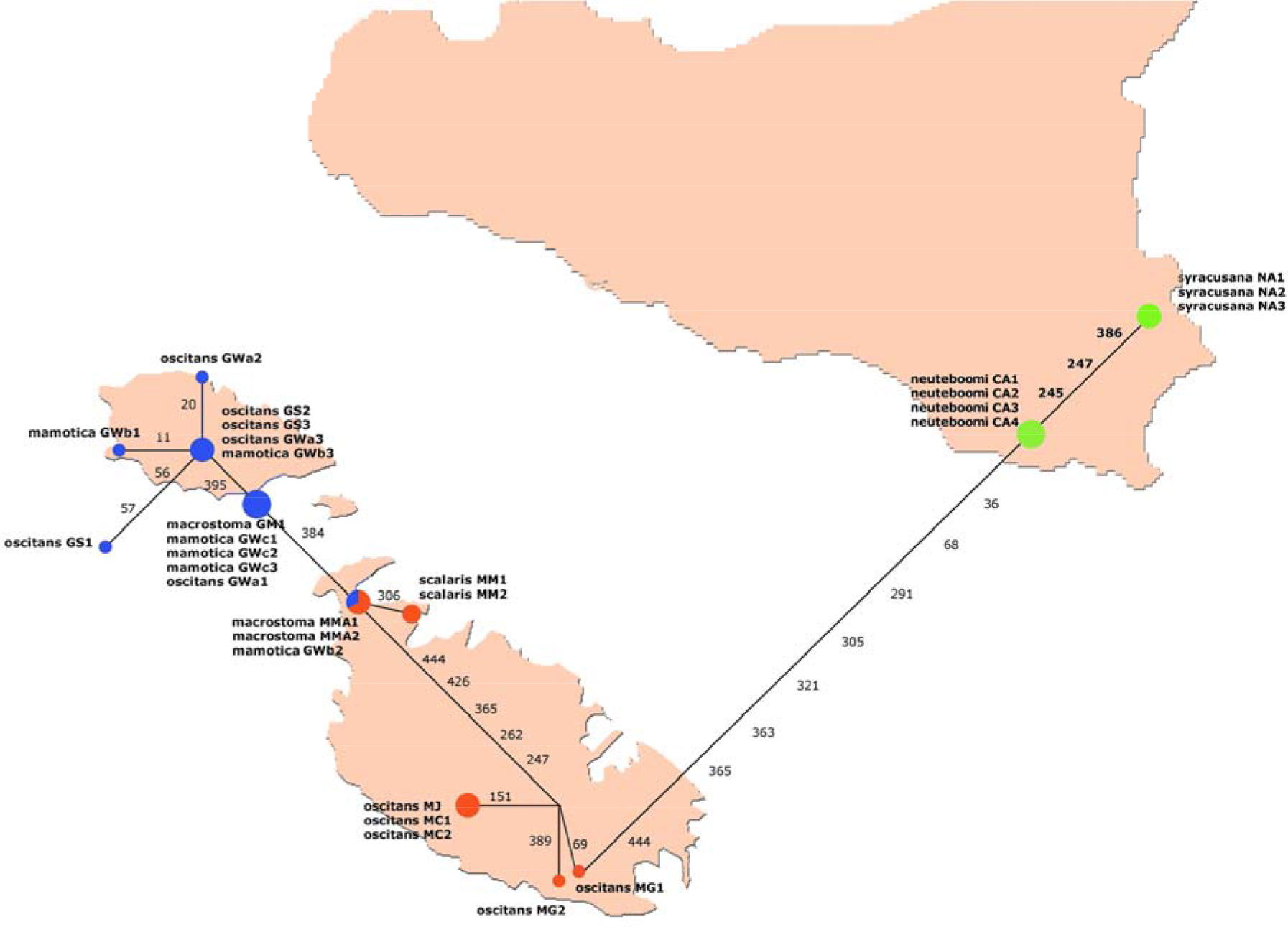
Median-joining network based on nuclear ITS-1 sequences across the *Muticaria* specimens. Numbers represent variable positions at sequences of the studied species. Diameter of the circles is directly proportional to the frequency and colours refer to geographical origin of the specimens (Red: Malta, Blue: Gozo, Green: Sicily). Codes are as in Table 1.

### MITOCHONDRIAL SEQUENCE VARIATION

As for *Lampedusa*, haplotype sequence divergence (HKY distances) between *L. lopadusae* and the population on Lampione islet was on average 0.03. Higher values were observed between *L. melitensis* and *L. imitatrix* from Malta, on average 0.12. Divergence between the Maltese *L. imitatrix* and *L. melitensis*, and *L. lopadusae*, was on average 0.20 and 0.22, respectively (similar values with the population from Lampione Islet: 0.23 and 0.24).

As for *Muticaria*, haplotype sequence divergence between *M. siracusana* and *M. neuteboomi* was on average 0.05. Divergences between Maltese *Muticaria* were on average: 0.07 between *macrostoma* and *oscitans*; 0.07 between *macrostoma* and *scalaris*; and 0.068 between *oscitans* and *scalaris*. Divergences between *Muticaria* from Gozo were on average: 0.05 between *macrostoma* and *oscitans*, and 0.03 between *oscitans* and *mamotica*.

## Discussion

### PHYLOGEOGRAPHIC RELATIONSHIPS

Phylogenetic analyses of the mitochondrial data (Fig. 8) clearly revealed that two well distinct evolutionary lineages occur across the Sicily Channel corresponding to the genera *Lampedusa* and *Muticaria* and that three parallel radiations might have occurred in the Maltese archipelago, one involving *Lampedusa*, one the Gozitan *Muticaria*, and the other the Maltese *Muticaria*.

These geographically structured lineages suggest that vicariance events may have played a substantial role in the pattern of diversification in this geographic area. This is noticeable within the *Lampedusa* lineage: taxa from Malta (*L. imitatrix* and *L. melitensis*) were well distinct, morphologically and genetically, from those from Lampedusa and Lampione (*L. lopadusae* and *L. nodulosa* or *L. l. lopadusae* and *L. l. nodulosa*). The disjunct distribution, the relatively high degree of genetic divergence, and the morphological differences may support an ancient vicariant event for the *Lampedusa* radiation, linked to the separation of the two island complexes (Pelagian and Maltese groups).

Phylogenetic analyses revealed a geographical structure within the *Muticaria* lineage as well. The two Sicilian *Muticaria* (*siracusana* and *neuteboomi*), which occur in south-eastern Sicily, constituted a distinct, well supported clade. Despite geographic closeness of the two sites sampled (about 20 km), both mitochondrial and nuclear data revealed a clear genetic distinction, already evidenced by Colomba et al. (2010). These two populations also differ significantly in shell characters although for some characters *neuteboomi* was not significantly different from some Maltese *Muticaria*. The origin of the morphological and genetic divergence of the two south-eastern Sicilian *Muticaria* clades remains unclear. A similar pattern in south-eastern Sicily was also found in cyprinonodontid freshwater fishes (Ferrito et al., 2007). It is possible that the geomorphology of the area (a relatively high Cenozoic calcareous plateau deeply separated by incised valleys) could have promoted fragmentation and isolation of populations. In fact, due to the extremely low vagility and a neighbourhood population structure of land snails (Wright, 1946; Schilthuizen & Lombaerts, 1994; Fiorentino et al., 2009) rapid genetic and morphological differentiation, even in geographically close populations, is not uncommon (Goodacre 2002; uit De Weerd, Piel & Gittenberger, 2004; Kameda, Kawakita & Kato, 2007).

As for the Maltese *Muticaria*, both *scalaris* and *oscitans* resulted monophyletic, while *macrostoma* was separated in two lineages. The three morphotypes, as well as the intermediate *macrostoma-oscitans* (for which genetic data were not available), were morphologically distinct (*oscitans* by NR; *macrostoma* and *scalaris* by the ratio D/H). Since only few samples of *macrostoma* were available, a fine local sampling is required to study the two *macrostoma* lineages in more detail. As in Sicilian *Muticaria*, the Maltese *Muticaria* also show a pattern of fine morphological and genetic geographical differentiation. In fact, the presence of morphologically significant *macrostoma-oscitans* intermediates may indicate that Maltese *Muticaria* are structured in demes across the island at a very local scale. A future more exhaustive sampling could unravel morphological and genetic microgeographical variability (i.e. many distinct genetic lineages corresponding to different morphotypes or a parapatric pattern between *macrostoma* and *oscitans*, as already supposed by Holyoak, 1986; Giusti et al., 1995).

The three Gozitan *Muticaria* morphotypes (*macrostoma*, *mamotica* and *oscitans*) were subdivided in three lineages, but with clear evidence of mixing in three cases: one *mamotica* (GWC1) grouped with *macrostoma*; one *mamotica* (GWB1) grouped with *oscitans*; one *oscitans* (GWA2) grouped with *mamotica*. Morphological analyses supported also this pattern of intermixed clades of the two morphotypes in *mamotica* and *oscitans*. Thus, the *mamotica* - *oscitans* clades showed a wide range of morphological variability, from the *oscitans* morphotype to the *mamotica* morphotype, the causes of which are still unknown (isolation *vs*. selection). However, it is worth noting that the *oscitans* and *mamotica* samples, which were mixed together in the same clades, came from the same locality. Thus, the two morphotypes are sympatric but genetically not monophyletic. This implies that the *mamotica* morphotypes appeared more than once and may be an adaptation. Morphotype *macrostoma* was also grouped with one sample of *mamotica* according to genetic analysis. Unfortunately, scarcity of *macrostoma* samples does not allow us to clarify this pattern. In general, these results indicate a possible relatively recent differentiation of Gozitan *Muticaria* or repetitive secondary contacts between different morphotypes. The latter hypothesis should be further investigated.

Allopatric differentiation seems to be the main mechanism underlying the radiation of the clausiilids across the Sicily Channel, although the sequence of events leading to the spread of the group still remains unclear according to mitochondrial data. The two outgroups used in this study, *Medora albescens* and *Leucostigma candidescens*, were resolved as distantly related taxa, providing no useful information on the origin of the group. Mitochondrial data do not contain sufficient phylogenetic signal to unequivocally infer the radiation within the *Muticaria* lineage, due to low resolution at internal nodes (Bayesian Analysis) and contrasting results between Bayesian and Parsimony reconstructions (Fig. 9). Difficulties in defining deep phylogenetic relationships in a tree are generally related to the effects of early and rapid diversification. Cladogenetic events occurring in close proximity might result in a lack of univocal phylogenetic signals, independent of the marker used (Albertson et al., 1999). However, Median-Joining Network analysis on nuclear ITS-1 sequences across *Muticaria* specimens (Fig. 10) and Maximum Parsimony analysis (Fig. 9) showed a closer relationship between haplotypes of Malta and Gozo than the Sicilian ones. Moreover, haplotypes of Gozo are strictly related to those of Malta but not to Sicily. Thus, considering that alopiine clausiliids are a mainly SE Euro-Mediterranean group (Nordsieck, 2007), colonisation events must have occurred through Sicilian corridors towards Malta and Gozo.

The origin of the Gozitan radiation from Malta is confirmed by the analysis of the nuclear ITS1 sequences. In fact, two specimens of *macrostoma* from Malta (site MC) and one *mamotica* from Gozo (site GWb2) shared the same nuclear variant. The most common nuclear variant present on Gozo was only one mutational step away from that found at site MC on Malta. A similar pattern is generally recognized as due to either lineage sorting of ancestral polymorphisms, or to gene flow between different lineages, or a combination of the two, since the two mechanisms are not mutually exclusive (Donnelly et al., 2004; Emerson & Oromí, 2005). In fact, shared haplotypes are randomly maintained in certain populations through incomplete lineage sorting. Rapid and recent radiations are consistent with this scenario since there would have been a short time for sorting of ancestral haplotypes into the descendant taxa. The star-like topology of the nuclear network for sequences from Gozo and the sharing of the same nuclear variant between specimens from Malta and Gozo may support this explanation.

### TAXONOMIC IMPLICATIONS

One goal of this study was to determine if *Muticaria* morphotypes previously described as formal taxa represent distinct evolutionary units. In the case of *Muticaria* from Malta, we found evidence supporting the monophyly of morphologically defined taxa such as *macrostoma*, *oscitans* and *scalaris*. *Muticaria* from Gozo, instead, where resolved as polyphyletic. For example, Gozitan specimens sampled at site GW, corresponding morphologically to *oscitans* and *mamotica*, were grouped together in the same subclade.

Incongruence between molecular and morphological evidence is not uncommon for land snails and has been repeatedly found even within the clausiliids. Repetitive parallel evolution of shell structures has been described for the clausilial apparatus in species belonging to the genera *Albinaria*, *Isabellaria*, and *Sericata* (Van Moorsel, Dijkstra & Gittenberger, 2000; Uit de Weerd et al., 2004; Uit de Weerd & Gittenberger, 2005). Moreover, a study on *Albinaria* based on mitochondrial data showed that specimens with strikingly different shell morphology (ribbed, semi-ribbed or smooth) and traditionally considered as different subspecies, are characterized by identical or very similar nucleotide sequences (Douris et al., 2007).

The phylogenetic results emerging from the present study indicate the need for a taxonomic re-evaluation of the included taxa. This raises the question of defining species boundaries. Several papers published in recent years and based wholly or partly on the same 16S rDNA region indicated that there is considerable variation in genetic distances at intraspecific level. Within other pulmonate species maximal sequence divergences have been reported i.e. 6% in *Cepaea nemoralis* (Davison 2000), 14% in *Euhadra quaesita* (Watanabe & Chiba 2001), 10% in *Partula* spp. (Goodacre 2002), 4.9% in *Candidula unifasciata* (Pfenninger & Posada 2002), and 23% in *Arion subfuscus* (Pinceel et al. 2004). For clausiliids, intraspecific values generally lower than 10% have been reported for populations of *Albinaria* spp., while distances between well distinct congeneric species are in the range 11-18% (Douris et al. 1998). However, it is clear that defining species boundaries cannot be reduced to a simple value of sequence divergence and other more valuable factors such as population history, geographical distribution of lineages and the presence of isolating barriers between them, should be taken in consideration.

*L. imitatrix* and *L. melitensis* may be considered distinct species, since they co-occur (not sympatrically) on the same island, and are morphologically distinct evolutionary lineages with high levels of genetic divergence (about 11%). On the other hand, the population of Lampione islet may be classified as a geographic form within *L. lopadusae*. The population from Lampione islet, however, is of particular conservation interest. Lampione is a small islet 700 m long by 180 m wide, located about 17 km northwest of Lampedusa. The risk of extinction is presumably high for taxa limited to very small areas such as the clausiliid population on Lampione. We stress that this population represents an important pool of genetic diversity within the Pelagian *Lampedusa*, and would argue for formal taxonomic recognition of this population (as *L. nodulosa* or *L. l. nodulosa*).

Considering the *Muticaria* lineage, phylogenetic relationships and observed mtDNA genetic distances would suggest that a definitive assessment is still difficult to achieve. The two entities occurring on Sicily can be considered two distinct taxa as they are morphologically and genetically distinguishable: *M. neuteboomi* and *M. siracusana* or *M. s. neuteboomi* and *M. s. siracusana*. Maltese *Muticaria* could be subdivided into three taxa according to morphological and molecular data (clade support and genetic divergence): *M. macrostoma* or *M. m. macrostoma, M. oscitans* or *M. m. oscitans* and *M. scalaris* or *M. m scalaris*. Gozitan *Muticaria* could be considered a distinct polytypic species (its oldest available name is *Muticaria mamotica*) subdivided into subspecies showing a morphological range from *macrostoma*-like to *mamotica*-like and *oscitans* like.

The taxonomic setting has important implications for the conservation of the clausiliids of the Sicilian Channel since legislation protecting species is usually based on recognizable taxa and does not normally take into account particular populations or sub-populations.. The *IUCN Red List of Threatened Species* designates *Lampedusa melitensis* as ‘Critically Endangered ( B1+2c)’, *Lampedusa imitatrix* as ‘Vulnerable (D2)’ and *Muticaria macrostoma* as ‘Lower Risk/near threatened’ under its 1994 ‘IUCN Red List Categories and Criteria version 2.3’ based on assessments made by Schembri (1996). Accession of new member states, including Malta, to the European Union (EU) in 2004 resulted in amendments to the EU’s ‘*Council Directive 92/43/EEC of 21 May 1992 on the conservation of natural habitats and of wild fauna and flora’*, better known as the ‘Habitats Directive’, to include amongst many other species, *Lampedusa imitatrix* and *Lampedusa melitensis* in Annexes II and IV. Annex II lists “Animal and plant species of community interest whose conservation requires the designation of Special Areas of Conservation”, while Annex IV lists “Animal and plant species of community interest in need of strict protection”. In Annex II, *Lampedusa melitensis* is further designated a ‘priority species’. The IUCN does not list any Italian species of *Lampedusa* and *Muticaria* and neither are any included in the EU’s ‘Habitats Directive’.

Therefore, while the Maltese *Lampedusa* species are adequately protected by international legislation (and also national legislation, since the requirements of the ‘Habitats Directive’ have been transposed to Maltese legislation), none of the other species/subspecies/populations/sub-populations of *Lampedusa* and *Muticaria* are, even if the areas occupied by some of these genetically distinct entities are of a few tens to hundreds of square metres only (see Giusti et al., 1995). Without formal taxonomic designations, it would be difficult to extend international legal protection to some of the more threatened populations, such as the *Lampedusa* of Lampione islet, the *‘scalaris’* population of Malta and the *‘mamotica’* populations of Gozo. In the interim period until the formal taxonomy of these entities is worked out, one solution may be to designate the more important and circumscribed populations of conservation importance as ‘evolutionarily significant units’ (ESUs) sensu Waples (1991) or as ‘management units’ (MUs) sensu Moritz (1994). ESUs are defined as populations that are reproductively separate from other populations and have unique or different adaptations while MUs are sets of populations that are currently demographically independent and which need to be managed independently of other populations for conservation purposes. Green (2005) recommends recognizing ‘designatable units’ (DUs) where not all populations of a species have the same probability of extinction, and therefore need different management strategies. According to Green (2005), DUs must be distinguishable on the basis of some morphological, genetic or distributional element and must have differing conservation status. It can be argued that the *Lampedusa* of Lampione islet, the *‘scalaris’* population of Malta and the ‘m*amotica*’ populations of Gozo qualify as ESUs, MUs and DUs on these criteria, particularly the last, since in this case, populations designated as DUs need not be evolutionary units but are determined by ecology and conservation status (Green, 2005, COSEWIC, 2009).

As already underlined (see discussion), since Maltese and Gozitan *Muticaria* were undersampled, it cannot be ruled out that a more exhaustive sampling in the area could show more lineages or areas of secondary contacts (occurred in the past or still present) where different morphotypes met. Thus, any scenario on the evolution of these clausiliids must be proposed with caution, and tested by further research.

## Acknowledgements

Alan Deidun, Enrico Talenti, Joseph Debono, Rosario Grasso and Simone Cianfanelli helped in field sampling or collected specimens for this research. PJS wishes to thank the University of Malta Research Committee for financial support.

